# Macroscale Adipose Tissue from Cellular Aggregates: A Simplified Method of Mass Producing Cell-Cultured Fat for Food Applications

**DOI:** 10.1101/2022.06.08.495192

**Authors:** John SK Yuen, Michael K Saad, Ning Xiang, Brigid M Barrick, Hailey DiCindio, Chunmei Li, Sabrina W Zhang, Miriam Rittenberg, Emily T Lew, Glenn Leung, Jaymie A Pietropinto, David L Kaplan

## Abstract

We present a method of producing bulk cell-cultured fat tissue for food applications. Mass transport limitations (nutrients, oxygen, waste diffusion) of macroscale 3D tissue culture are circumvented by initially culturing murine or porcine adipocytes in 2D, after which bulk fat is produced by mechanically harvesting and aggregating the lipid-filled adipocytes into 3D fats using alginate or transglutaminase binders. The 3D fats were visually similar to fat tissue harvested from animals, with matching textures based on uniaxial compression tests. The mechanical properties of cultured fat tissues were based on binder choice and concentration, and changes in the fatty acid compositions of cellular triacylglyceride and phospholipids were observed after lipid supplementation (soybean oil) during *in vitro* culture. This approach of aggregating individual adipocytes into a bulk 3D tissue provides a scalable and versatile strategy to produce cultured fat tissue for food-related applications, thereby addressing a key obstacle in cultivated meat production.

**Graphical Abstract:** 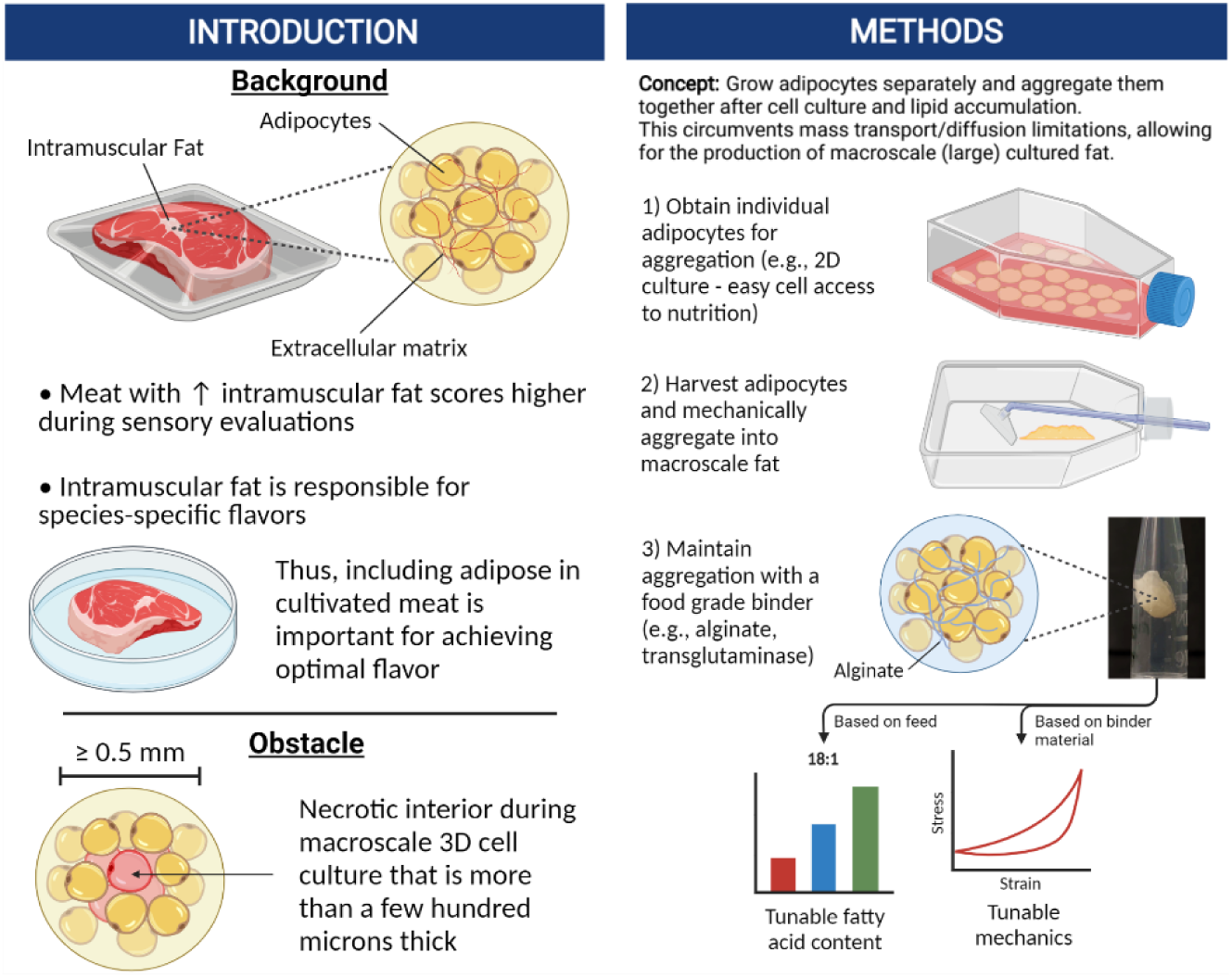

## Introduction

Cultivated meat (also cultured, cell-cultured, *in vitro* meat) uses tissue engineering to produce meat for food^1–3^. Canonically, this research has focused on recreating the muscle component of meat^2,4–8^. However, fat is key to the taste and texture of meat^9–12^. For example, peak evaluation scores were achieved with beef samples containing ∼36% crude fat^13^. Hundreds of volatile compounds are released when meat is cooked, with a majority originating from lipids^12,14,15^. Fat is also responsible for the species-specific flavor of meat, making it important for reproducing the flavors of specific animals^16^. In addition to cultivated meat, *in vitro*-grown fat could also be used to enhance existing plant-based meats, as complex species-specific flavors of animals are difficult to recapitulate *de novo*^17–19^.

While cultured fat stands to greatly improve the quality of alternative proteins, producing macroscale tissues remains a major challenge due to mass transport limitations. Even millimeter-scale *in vitro* tissues are challenging to generate, as oxygen and nutrients are only able to diffuse through several hundred microns of dense tissue^20–23^. Various approaches have been implemented in tissue engineering to overcome diffusion limitations, such as vascularization and the incorporation of perfusion channels^24–34^. However, current strategies are often limited by scalability and complexity. A dense array is required when incorporating perfusion channels, as all tissue must remain within several hundred microns of a channel^35^. For adipose tissue, lipid accumulation decreases considerably once cells are ∼1 mm away from a perfusion channel^26^. Vascularization of macroscale tissues is achieved via co-culture with endothelial cells, which spontaneously organize into capillary-sized vessels^36^. However, spontaneous vessel formation may not be feasible for centimeter to meter-scale tissues, as cultivated cells may die during the time required for the vasculature to form^37^. Capillaries may also be too small to deliver a sufficient volume of media when supporting larger tissues^38,39^. Thus, many challenges remain to scale fat (and other) tissues to bulk production levels.

In this study, our goal was to develop a relatively simple method of producing bulk cultured fat that circumvents contemporary mass transport limitations. This was achieved by growing murine and porcine adipocytes in thin layers with easy access to the culture media, followed by post-growth aggregation into 3D adipose after sufficient adipocyte maturation. Aggregation at the end of cell culture removed the need for nutrient delivery via vascularization or elaborate 3D tissue perfusion, thus reducing costs and improving scalability. This approach is feasible when creating tissue solely for food purposes, as there is no requirement for continued cell survival once the final edible tissue is produced. This paradigm allows for simpler approaches and outcomes in comparison to regenerative medicine goals.

We hypothesized that aggregating individual adipocytes would be sufficient to reproduce the taste, nutrition, and texture profile of fat, as adipose tissue *in vivo* is a dense aggregation of lipid filled adipocytes with a sparse extracellular matrix (ECM)^40^. Lipid-filled adipocytes were grown *in vitro* and aggregated into bulk macroscale tissues within suitable matrices, then compared with native adipose tissue from various animals. Cultured fat and native adipose samples were characterized for lipid droplet morphology and mechanical properties to infer textural characteristics. The fatty acid compositions of cultured adipocytes were also analyzed to garner insight into flavor and nutrition.

## Materials and Methods

### Preadipocyte Cell Culture

Adipogenic mouse (*Mus musculus*) 3T3-L1 cells were obtained from the American Type Culture Collection (CL-173; ATCC, Manassas, VA, USA) and used to generate cultured fat tissues. 3T3-L1s were received at passage X+13 (with X being an unknown number of passages as stated by ATCC) and expanded in high glucose Dulbecco’s modified eagle medium (DMEM) with GlutaMAX, phenol red and sodium pyruvate (10569044; ThermoFisher, Waltham, MA, USA) supplemented with 10% bovine calf serum (12133C; Sigma, Burlington, MA, USA) and 1% antibiotic/antimycotic (Anti/Anti, 15240062; ThermoFisher) in a 37°C, 5% CO_2_ incubator. General cell counting was performed using an automated cell counter (NucleoCounter NC-200; Chemometec, Lillerød, Denmark) and cell detachment was achieved enzymatically using Accumax (AM105; Innovative Cell Technologies Inc, San Diego, CA, USA). For cryopreservation, the same medium was mixed with dimethylsulfoxide (DMSO, D2438; Sigma) at a ratio of 9:1 (medium:DMSO) and cells were frozen overnight at -80°C in an isopropanol-based freezing container (5100-0001; ThermoFisher). (Long term storage in liquid nitrogen) 3T3-L1s were used between passages X+15 and X+18.

Dedifferentiated (DFAT) porcine (*Sus domesticus*) cells were generated by isolating mature adipocytes from the subcutaneous adipose tissue of a Yorkshire pig (93 days of age) and was based off several studies on porcine DFAT cells in the literature^41,42^. In brief, porcine tissue was procured from the Tufts University Medical Center and transported to the laboratory on ice (∼25 minutes). The tissue was disinfected with a solution of Dulbecco’s phosphate buffered saline (14190250; ThermoFisher) containing 10% Anti/Anti. 5 g of tissue was collected from the disinfected tissue and minced into <1 mm^3^ pieces in the DPBS-Anti/Anti solution. The tissue was then incubated in 25 ml of DMEM containing 0.1% collagenase (LS004176; Worthington Biochemical, Lakewood, NJ, USA) and 10% Anti/Anti for 1.5 hours at 37°C, then 45 minutes at room temperature (RT). During incubation, the collagenase-adipose solution was inverted every 15 minutes. Afterwards, the collagenase-adipose solution was filtered through a 300 μm cell strainer (43-50300-03; pluriSelect, San Diego, CA, USA) and centrifuged for 5 minutes at 500 x g. Mature adipocytes were collected from the floating lipid layer in the centrifuged solution and added to 10 ml of DMEM containing 20% fetal bovine serum (FBS, A31606-01, Lot: 2129571; ThermoFisher) and 100 μg/ml Primocin (ant-pm-1; InvivoGen, San Diego, CA, USA), then centrifuged again (500 x g, 5 minutes). Mature adipocytes were collected and transferred to a T25 tissue culture flask (Nunc EasYFlask, 156367; ThermoFisher) containing 3.5 ml of DMEM with 20% FBS and 100 μg/ml Primocin and incubated at 39°C with 5% CO_2_. After 2 days, the floating lipid layer was transferred into a new T25 flask and completely filled (∼75 ml) with DMEM + 20% FBS + 100 μg/ml Primocin. The filled flask was kept at upside down at 39°C, 5% CO_2_ for 5 days, during which the mature adipocytes attached to the top surface and dedifferentiated. The flask was then drained, flipped around, and used normally with 3.5 ml of culture media. DFAT cells were used up to passage 3, grown using DMEM + 20% FBS + 100 μg/ml Primocin, and cryopreserved with a 9:1 solution of culture medium and DMSO.

### Adipogenic Differentiation

During terminal passages (where cells were differentiated instead of passaged for further cell culture), 3T3-L1s were grown DMEM with 10% FBS and 1% Anti/Anti until 100% confluency. 24-48 hours after reaching confluency, cells were switched to an adipogenic induction medium (differentiation medium) consisting of DMEM with 10% FBS, 1% Anti/Anti, 10 μg/ml insulin (I0516; Sigma), 1 μM dexamethasone (D4902; Sigma), 0.5 mM 3-isobutyl-1-methylxanthine (IBMX, I5879; Sigma) and 2 μM rosiglitazone (R0106; TCI America, Portland, OR, USA). After 2 days of adipogenic induction, 3T3-L1s were switched to a lipid accumulation medium for 13-28 days with feeding every 2-3 days. During cell feedings, great care was taken to handle the cells gently to prevent lift-off of lipid-laden adipocytes. The lipid accumulation medium consisted of DMEM with 10% FBS and 1% Anti/Anti, with 120 μg/ml Intralipid (I141; Sigma) and 10 μg/ml insulin. For experiments on the effects of fatty acid supplementation on cultured adipocytes, Intralipid concentration varied from 0-1000 μg/ml. “15-day” cultured adipocytes refer to 2 days of adipogenic induction and 13 days of lipid accumulation post-induction, while “30-day” adipocytes refer to 2 days of induction and 28 days of accumulation.

Porcine DFAT cells were differentiated similarly to 3T3-L1s, except the adipogenic induction and lipid accumulation phases lasted for 4 and 26 days respectively (total 30 days). Different adipogenic media were also used. The induction medium consisted of Advanced DMEM/F12 (12634-028; ThermoFisher) supplemented with 2 or 20% FBS, 0 or 500 μg/ml Intralipid, 100 μg/ml Primocin, 0.5 μM dexamethasone, 0.5 mM IBMX, 5 μM rosiglitazone, 2 mM (1X) GlutaMAX (35050-061; ThermoFisher), 20 μM biotin (B04631G; TCI America, Portland, OR, USA), and 10 μM calcium-D-pantothenate (P001225G; TCI America). During lipid accumulation, the same medium was used sans IBMX.

### Harvest of Lipid-Laden Adipocytes and Formation of 3D Cultured Fat Constructs

After adipogenic differentiation, adipocytes were rinsed with DPBS after gentle aspiration of the culture media. Flasks were kept on their sides for 5-10 minutes to more thoroughly drain the DPBS, then cells were harvested using a cell scraper via a series of raking motions (as opposed to the windshield wiper approach) into a pre-weighed 50 ml centrifuge tube.

All reconstructed cultured fat tissues were generated in cylindrical 3D printed molds open from the top and bottom (Ø 6 mm, 3 mm height) using a biocompatible resin (Surgical Guide resin; Formlabs, Somerville, MA, USA) and a stereolithography 3D printer (Form2 Printer; Formlabs). A glass cover slip was adhered (Loctite 3556; Henkel Adhesives, Düsseldorf, Germany) to the bottom of the mold to hold the adipocytes during tissue formation and was taken off afterwards to facilitate tissue removal. Two approaches were carried out to form the 3D cultured fat tissues. The first method involved mixing cell scraped *in vitro* adipose with a sterile-filtered 15% (w/v) microbial transglutaminase (mTG) solution (Activa TI; Ajinomoto, Itasca, IL, USA) at a 4:1 ratio (adipose:transglutaminase) and incubating the mixed tissue at 37°C overnight. The second method involved embedding the *in vitro* adipose in slow gelling alginate of various concentrations, with the approach adapted from the literature^43^. Here, 1.6% or 3.2% (w/v) medium viscosity sodium alginate solutions (1007-50; Modernist Pantry, Eliot, ME, USA) were prepared by adding powder to water and allowing the alginate to hydrate for at least 1 hour. After hydration, calcium carbonate powder (CaCO_3_, 1505-50; Modernist Pantry, Eliot, ME, USA) was added to the alginate solutions to achieve 25 mM and vortexed for 10-60 seconds to yield evenly dispersed suspensions. To initiate delayed gelation, glucono delta-lactone powder (1159-50; Modernist Pantry, Eliot, ME, USA) was added to the alginate-calcium mixtures to achieve 50 mM and vortexed again for 10-60 seconds. Finally, the 1.6% and 3.2% alginate solutions were mixed 1:1 (volumetrically) with cell scraped adipocytes to obtain alginate-cultured fat constructs of 0.8% and 1.6% alginate respectively. For cultured porcine fat, scraped adipocytes were mixed with 3.2% alginate at a 1:1 (volumetric) ratio. All macroscale cultured fat tissues were stored at 4°C for a maximum of 3 days prior to mechanical testing.

### Native Animal Adipose Samples

Beef fat (*Bos taurus*) from the pectoralis minor (brisket point cut) and chicken fat (*Gallus gallus*) from the thigh were purchased from a local butcher shop. Both samples were displayed at refrigerator temperatures in the shop and transported to the laboratory at RT (∼15 minutes). Two types of pork fat were used in this study. For lipidomics, pork fat from the belly was purchased from a local butcher shop. For mechanical testing and fluorescence imaging, adipose from the belly of a freshly sacrificed 97-day old Yorkshire pig was obtained at the Tufts University Medical Center and transported to the laboratory on ice (∼25 minutes). Food-grade lard (Goya, Secaucus, NJ, USA) and tallow (Epic Provisions, Austin, TX, USA) were purchased commercially for mechanical testing and shaped with the same 3D printed molds used for forming macroscale *in vitro* fats. For lipidomics, beef, pork and chicken samples were flash frozen in liquid nitrogen and stored at -80°C, then thawed for lipid extraction. For mechanical (compressive) testing and fluorescence imaging, beef, pork and chicken adipose samples were directly used after acquisition. Mouse fat samples were obtained from the perigonadal regions of 7- and 67-day old mice (CD-1 strain; Charles River, Wilmington, MA, USA) and directly used for lipidomics and fluorescence imaging.

### Mechanical Testing

Unconfined uniaxial compressive testing was carried out using a dynamic mechanical analyzer (RSA3; TA Instruments, New Castle, DE, USA). Cultured fat samples (Ø 6 mm, 3 mm height) were taken from storage at 4°C and allowed to warm to RT (25°C as measured by an infrared thermometer) prior to testing. Beef, pork, and chicken fat samples were cut using 6 mm biopsy punches, cut to 3 mm heights and tested at 25°C. During testing, fat samples were placed between parallel stainless steel plates and compressed. Compressive stress responses and elastic recoveries were measured up to 50% strain at a rate of 1.667% s^-1^. Stress/strain curve data were obtained using the TA Universal Analysis software.

### Fluorescence and Immunofluorescence Staining

Cultured adipocytes in 48 well tissue culture plates were stained to confirm lipid accumulation and to check for markers of adipogenic differentiation. Here, 15-day and 30-day 3T3-L1s adipocytes were stained for neutral lipids using DMEM 10% FBS 1% Anti/Anti culture media supplemented with 2 μM BODIPY 493/503 (4,4-difluoro-4-bora-3a,4a-diaza-s-indacene, D3922; Invitrogen, Waltham, MA, USA). Cells were first rinsed with DPBS, then incubated at 37°C for 15-30 minutes with 140 μl of the BODIPY-culture media solution in each well. After BODIPY incubation, cells were rinsed twice with DPBS and then fixed in 4% PFA for 10-15 minutes at RT protected from light. After fixation, cells were rinsed twice in DPBS then simultaneously blocked and permeabilized with B/P buffer [DPBS with 0.1% Tween 20 (P2287; Sigma), 0.1M glycine (161-0718; Bio-Rad, Hercules, CA, USA), 0.02% sodium azide (S2002; Sigma) and 5% goat serum (16210-064; ThermoFisher)]. Cells were then incubated overnight with primary antibodies at 4°C in B/P buffer containing either 1:50 rabbit anti-fatty acid synthetase (MA5-14887; Invitrogen, Waltham, MA, USA) or 1:500 rabbit anti-peroxisome proliferator-activated receptor gamma (ab45036, Abcam, Waltham, MA, USA). The next day, cells were rinsed in DPBS and incubated with 2 μg/ml DAPI (4′,6-diamidino-2-phenylindole, 62248; ThermoFisher) and 1:500 Alexa Fluor Plus 647 conjugated goat anti-rabbit secondary antibodies (A32733; Invitrogen, Waltham, MA, USA) for 1 hour at RT. Finally, cells were rinsed in DPBS twice and stored in mounting media (H-1700; Vector Laboratories, Burlingame, CA, USA) at RT. Porcine DFAT adipocytes were stained for lipids via a 15-30 minute incubation at 37°C in lipid accumulation media (sans Intralipid) supplemented with 2 μM BODIPY 493/503 and 2 μg/ml Hoescht 33342 (H3570; Invitrogen). DFAT cells were then fixed in 4% PFA at RT for 10-15 minutes, then rinsed with DPBS and stored in mounting media.

For lipid staining of native adipose samples, chicken, beef, pork, and mouse tissues were first fixed in 4% PFA at 4°C overnight. After fixation, adipose tissues were incubated in optimal cutting temperature compound (OCT, 4583; Sakura Finetek, Torrance, CA, USA) overnight at 4°C. Samples were then placed in molds with additional OCT and flash frozen with liquid nitrogen. Blocks of fat embedded OCT were then cryosectioned (Leica CM1950; Wetzlar, Germany) and dried on glass slides (Superfrost Plus Gold, 22-035813; Fisher Scientific, Waltham, MA, USA) at RT for 15-30 minutes. After drying, slides with tissue sections were rehydrated in DPBS for 10 minutes to dissolve the OCT, then stained with 2 μM BODIPY 493/503 for 30 minutes at RT. Stained tissues were then cleaned with DPBS and mounted with a coverslip plus mounting media. Staining jars (EasyDip; Simport Scientific, Saint-Mathie-de-Beloeil, QC, Canada) were used whenever possible to minimize sample loss that was sometimes observed when pipetting solutions directly onto the glass slides.

Z-stack images of all samples were taken using a confocal microscope (SP8; Leica, Wetzlar, Germany) using 405, 488, 638 nm lasers as appropriate. Quantifications of lipid accumulation (degree of BODIPY staining) were performed by first segmenting images into stain versus background regions using ilastik, followed by measurements of the stain area in CellProfiler^44,45^. BODIPY staining area was divided by the number of nuclei detected to calculate the amount of lipid accumulation per cell. PPARγ % was calculated by counting the number of PPARγ positive nuclei in CellProfiler (after ilastik segmentation), then dividing by total nuclei. Lipid droplet diameters were measured by first segmenting BODIPY images in ilastik, then taking the mean diameter of each BODIPY object (lipid droplet) in CellProfiler.

### Lipid Extraction

Lipids were extracted from *in vitro* adipocytes and native adipose tissues for downstream lipidomics analyses. For this analysis, *in vitro* adipocytes were rinsed three times with DPBS with 5 minute waits to remove residual culture media, followed by detachment with a cell scraper after the removal of residual DPBS via vacuum aspiration. This cell scraped cultured fat was used for lipid extractions and lipidomics. Native adipose samples were thawed from -80°C storage (previously flash frozen with liquid nitrogen) and 10 mg (pork, beef, chicken) or 15 mg (mouse) samples were minced with tweezers and scissors prior to lipid extraction. All groups were performed in triplicate.

Lipid extractions were performed using a methyl tert-butyl ether-based (MTBE, 41839AK; Alfa Aesar, Haverhill, MA, USA) approach scaled down for 1.5 ml Eppendorf tubes^46^. 400 μl of methanol (BPA4544; ThermoFisher) was added to the fat samples in a 1.5 ml centrifuge tube and vortexed for 1 minute, followed by an addition of 500 μl MTBE with occasional mixing for 1 hour at RT. Next, 500 μl of water was added and solutions were vortexed for 1 minute. Aqueous and organic phases were separated by centrifuging samples at 10,000 x g for 5 minutes. The upper organic phase was then transferred to a new tube while the aqueous phase was re-extracted using 200 μl of MTBE, 1 minute of vortexing and 5 minutes of centrifugation. The combined organic phases were then dried under nitrogen and stored at - 80°C.

### Lipidomics Analyses

Lipidomics analyses were performed at the Beth Israel Deaconess Medical Center Mass Spectrometry Facility (John Asara Laboratory) according to prior procedures^47^. In brief, liquid chromatography-mass spectrometry/mass spectrometry (LC-MS/MS) using a ThermoFisher QExactive Orbitrap Plus mass spectrometer coupled with an Agilent 1200 high pressure liquid chromatography (HPLC) system and an Imtakt Cadenza CD-C18 HPLC column (CD025, 3.0 μm particle size, 2.0 mm inner diameter x 150 mm length) was used to characterize lipid samples. Lipid species identification and peak alignment was achieved using LipidSearch (ThermoFisher), after which the prevalence of different fatty acids within triacylglycerides and phospholipids were quantified using a custom Python script which is available at: https://github.com/mksaad28/Yuen_et_al.

### Statistical Analyses

Statistical analyses were carried out in GraphPad Prism 9.3.0. The specific analyses used are indicated in the text and include: Analysis of variance (ANOVA) with Tukey’s post-hoc tests, t-tests with Welsh’s correction, principal component analyses, and chi-square tests. Error bars and ± ranges represent standard deviations unless specified with SEM (standard error of the mean). 0.05 was the cut-off P value for statistical significance. All experiments in this study were carried out with at least triplicate samples (n ≥ 3). The aggregation of 2D grown adipocytes into bulk 3D tissues has been repeated over three times over numerous experiments.

## Results

### Murine adipogenic precursor cells accumulate lipid and express adipogenic markers during 2D *in vitro* culture, resembling native adipocytes after 30 days of adipogenesis

To obtain individual fat cells needed to test the concept of adipocyte aggregation into 3D fat tissue, we cultured murine 3T3-L1 cells using 2D culture for good cellular access to nutrition and simple cell harvest. After reaching confluency, 3T3-L1s were induced to differentiate into adipocytes for 2 days, then switched to lipid accumulation media until 15-30 days of culture (**Figure 1A**). At day 15, cultured adipocytes were 82.5% (± 5.2%) positive for peroxisome proliferator-activated receptor gamma (PPARγ) (**Figure 1B, Supplemental Table 1**). 30-day adipocytes contained ∼3 times more lipid per cell than 15-day adipocytes (**Figure 1C,D**). During cell culture, gentle treatment of the cells (e.g., when changing culture media) was key to preventing lift-off of lipid-filled adipocytes (data not shown). A positive correlation was observed between adipogenic culture duration and lipid droplet size (**Figure 1E, Supplemental Table 2**). Mean lipid droplet diameters for 15- and 30-day adipocytes were 8.1 μm (± 0.04 SEM) and 11.1 μm (± 0.1 SEM) respectively. Cultured adipocytes exhibited the appearance of packed lipid droplets observed when imaging native adipocytes from various animals, particularly that of 7-day old mouse (P7 mouse) and chicken (**Figure 1F**).

**Figure 1.**
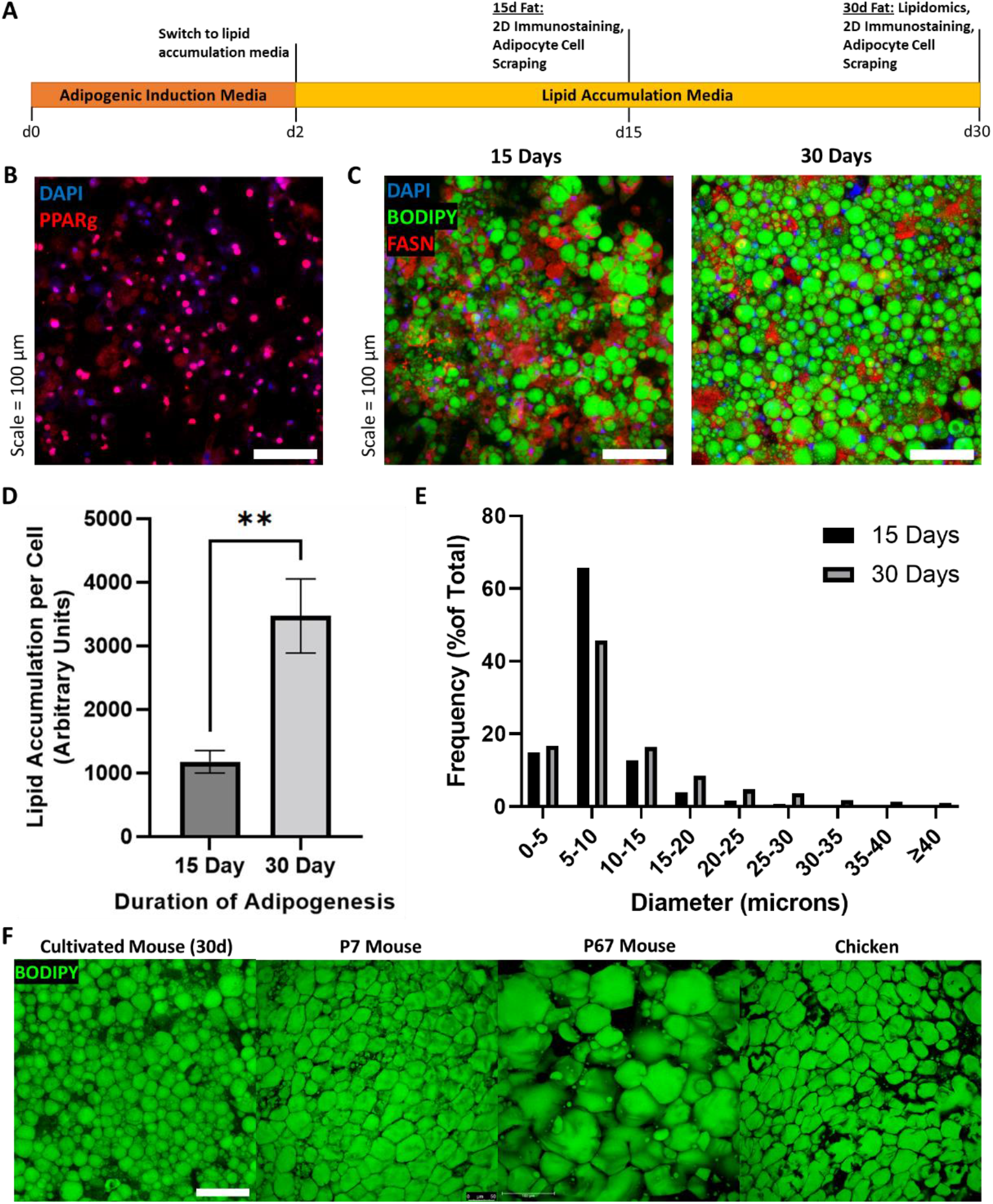
**(A)** Timeline of 3T3-L1 adipogenic differentiation. Confluent preadipocytes were grown in adipogenic induction medium for 2 days, then switched to lipid accumulation media for 15 or 30 days, where cells were stained and imaged, or harvested for lipidomics and 3D cultured fat tissue formation. **(B)** 15-day adipocytes stained for the adipogenic transcription factor PPARγ (red), as well as nuclei via DAPI (blue). **(C)** Lipid stained (BODIPY, green) adipocytes after 15 and 30 days of adipogenesis. The *in vitro* adipocytes were also stained for DNA via DAPI (blue) and fatty acid synthetase (red). **(D)** The mean degree of lipid accumulation in 15- and 30-day cultured adipocytes, normalized by the number of cells detected via nucleus counting. n = 4 for each sample group compared using an unpaired t test with Welch’s correction, where p ≤ 0.01 (**). **(E)** Frequency distributions of lipid droplet diameters from cultured adipocytes adipogenically grown for 15 and 30 days. n = 4 for each sample group, compared via chi-square test in **Supplemental Table 2. (F)** Lipid staining images (BODIPY) of 30-day *in vitro* 3T3-L1s compared to native adipocytes from chicken and mice of two ages. “P7 Mouse” and “P67 Mouse” refer to 7- and 67-day old mice respectively. Scale bar (same for all images) represents 100 μm.

### Aggregated *in vitro* adipocytes form macroscale cultured fat constructs

After 15-30 days of adipogenesis, cultured adipocytes were harvested with a cell scraper **(Figure 2A, B**). Adipocytes were scraped mechanically due to insufficient detachment when using Accumax (incubation time: >20 minutes, data not shown). During cell harvest, cultured adipocytes aggregated by the mechanical action of the cell scraper appeared like fat tissue or lipoaspirate (**Supplemental Figure 1**).We found that one culture flask with 175 cm^2^ surface area generally yielded ∼0.8 g of cell scraped adipocytes (data not shown). After harvest, scraped adipocytes successfully formed macroscale 3D tissues that subsequently stayed as one-mass after mixing with binders to hold individual cells together (**Figure 2C,D**). Either microbial transglutaminase (mTG) or alginate were used as the fat binders (**Supplemental Figure 2**).

**Figure 2.**
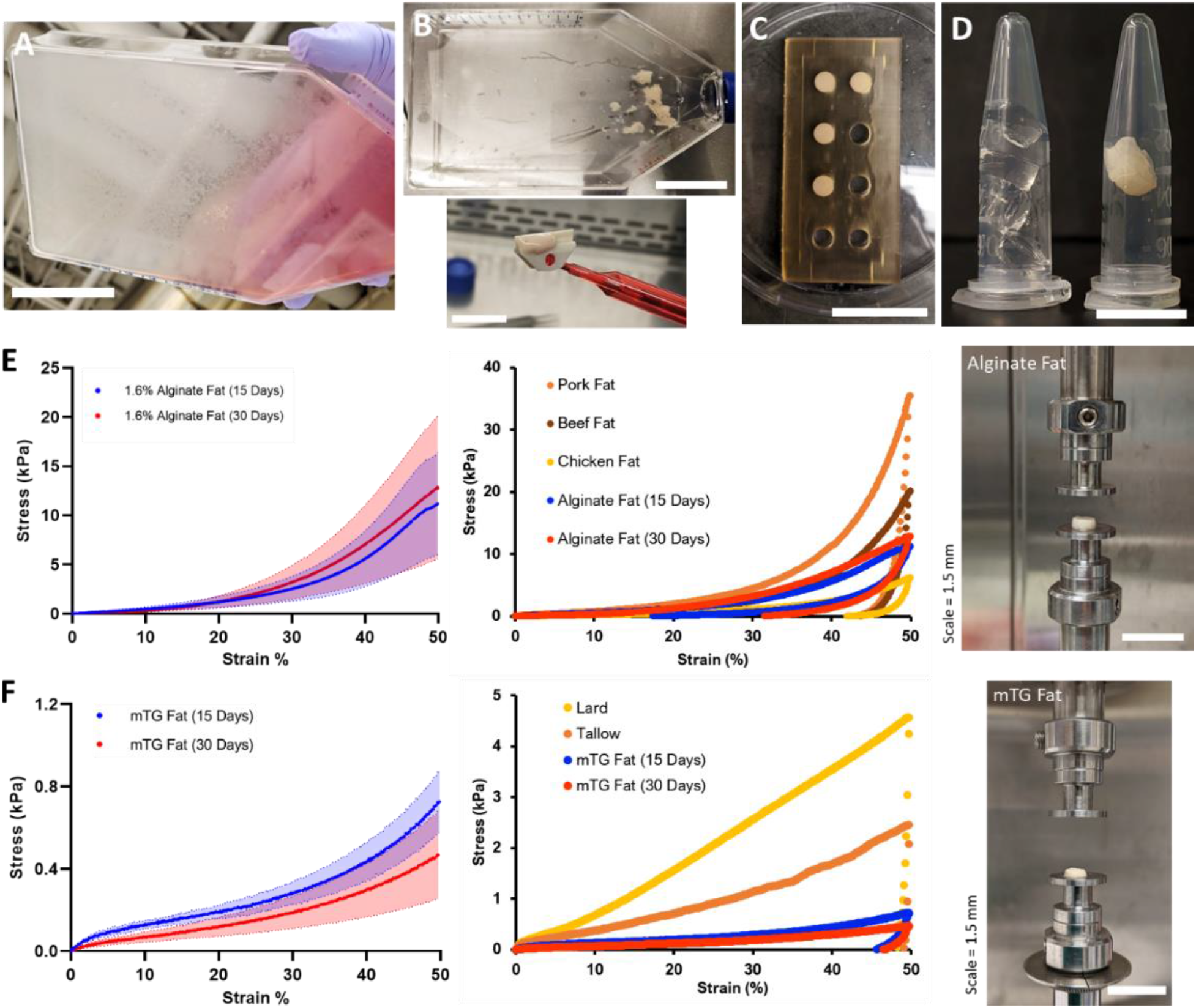
**(A-D)** Steps for producing 3D macroscale cultured fat through the aggregation of individual adipocytes grown *in vitro*. **(A)** <u>Adipogenesis:</u> Adipocytes were differentiated *in vitro* for 15 to 30 days. Lipid-accumulating fat cells turned the bottom of the cell culture flask opaque. Scale bar 5 cm. **(B)**<u>Cell Harvest:</u> Adipocytes were mechanically collected with a cell scraper, which also aggregated the cells into masses of cultured fat. Scale bars 5 cm and 1.5 cm for the top and bottom images respectively. **(C)**<u>Binding into 3D Tissue:</u> Harvested fat was combined with a binder (e.g., alginate, transglutaminase) in a mold to add structure. Scale bar 3 cm. **(D)** <u>3D Cultured Fat:</u> Cylinders of structured cultured fat after removal from the mold. 3D cultured fat made with 1.6% alginate is shown in the right tube, while the left tube contains 1.6% alginate without fat cells. Scale bar 1 cm. **(E and F)** Mechanical testing of cultured and native fat tissues (uniaxial compression). **(E and F, Left)** The compressive strength of alginate-based (E) and microbial transglutaminase (mTG)-based (F) cultured fats with 15 or 30-day adipocytes. The solid lines represent mean values, while the shaded areas represent standard deviations. **(E, Middle)** A compressive strength comparison of alginate-based cultured fats with intact pig, cow and chicken adipose tissues. Data points represent mean values, and the overall data represent tissue loading from 0 to 50% strain over 30 seconds, followed by unloading to 0% strain over the same duration. **(F, Middle)** A compressive strength comparison of mTG fats with rendered animal fats from pigs (lard) and cows (tallow). Data points represent mean values, and the overall data represents tissue loading from 0 to 50% strain over 30 seconds, followed by unloading to 0% strain over the same duration. **(E and F, Right)** The appearance of macroscale alginate (E) and mTG (F) cultured fats on the testing apparatus prior to compression.

### Mechanical properties of macroscale cultured fats

After forming macroscale constructs, cultured fat tissue samples were subjected to uniaxial compression testing as a proxy for food texture. Sodium alginate and mTG as “generally recognized as safe” (GRAS) materials in the United States were selected as binders for cultured adipocytes, with precedent for alginate as a fat replacer and mTG as a binder in meat products^48,49^. For alginate-fats, 1.6% (final concentration) was selected after preliminary experiments (**Supplemental Figure 3**). The degree of adipocyte lipid accumulation (cultured fat tissues produced using 15-versus 30-day old adipocytes) did not have a significant impact on resistance to compression (**Figure 2E and F, Left**). Alginate fats containing 15- and 30-day adipocytes exhibited mean compressive stresses of 11.1 kPa (± 5.2) and 12.8 kPa (± 7.3) at 50% strain respectively (p = 0.7304, Welch’s t-test). For 15- and 30-day mTG fats, the mean resistances to compression were 0.5 kPa (± 0.2) and 0.7 (± 0.2) at 50% strain respectively (p = 0.1030, Welch’s t-test) (**Figure 2F, Left**).

**Figure 3.**
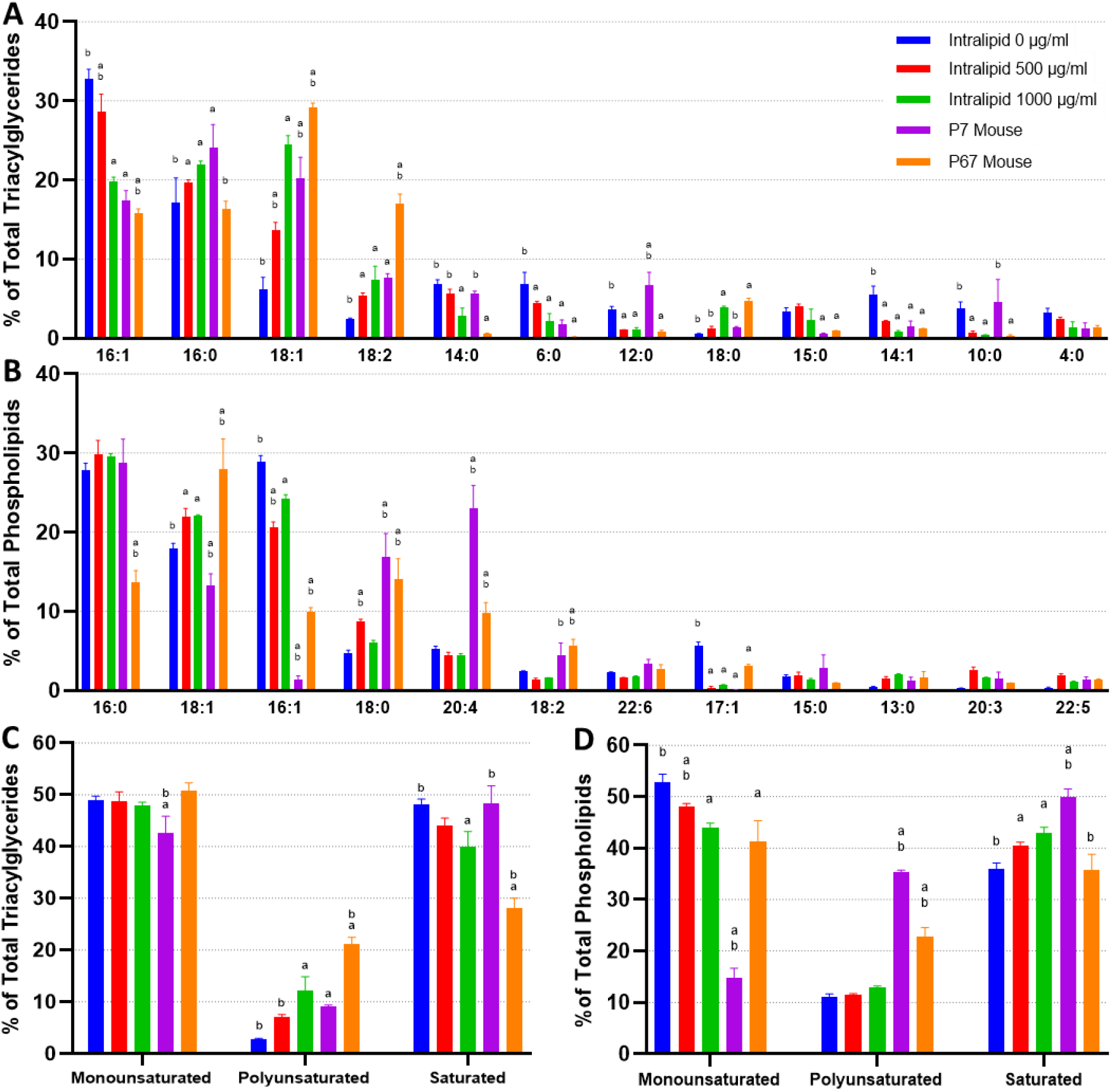
The fatty acid composition of **(A)** <u>triacylglycerides</u> and **(B)** phospholipids from *in vitro* (30 days of adipogenesis, 0-1000 μg/ml Intralipid) and *in vivo* <u>murine</u> fats. The top 12 most prevalent fatty acids across all the sample groups are shown. Tables containing the proportions of all detected fatty acids, as well as PCA graphs, are available (**Supplemental Tables 4 and 5, Supplemental Figure 5**). **(C)** and **(D)** represent fatty acids categorized by degree of saturation for triacylglycerides and phospholipids respectively. The top-down sample order in the legend is the same as the left-right order of the samples in the column charts. ‘a’ and ‘b’ represent a difference of p ≤ 0.05 versus 0 μg/ml Intralipid and 1,000 μg/ml Intralipid respectively.

The stress-strain profile of alginate-based cultured fats were within the general range of livestock and poultry fat tissues, while displaying similar hysteresis behavior during loading and unloading (**Figure 2E, Middle**). At 50% strain, mean compressive stresses for beef and pork fat were 20.2 kPa (± 12.5) and 35.6 kPa (± 17.5), while the mean value for chicken fat was 6.2 kPa (± 2.0). mTG-based cultured fats were more similar to rendered fats, although lard and tallow samples appeared stronger with mean compressive stresses of 4.6 kPa (± 1.4) and 2.5 kPa (± 0.9) at 50% strain respectively (**Figure 2F, Middle**). mTG-based cultured fats maintained some resistance during unloading, while rendered fats did not rebound after compression.

### The fatty acid compositions of *in vitro* murine adipocytes can be tuned by lipid supplementation during cell culture

Next, we looked to compare the intracellular lipids of *in vitro* and *in vivo*-grown mouse adipocytes. *In vitro* adipocytes were additionally cultured with 0 – 1,000 μg/ml of Intralipid (a soybean oil-based lipid emulsion) to investigate the potential effects of fatty acid (FA) supplementation on adipocyte lipids.

For *in vitro* adipocytes (also termed *in vitro* fats, IVFs) without lipid supplementation (other than lipids in FBS), major differences in intracellular triacylglycerides (TAGs) versus native mouse fats included an enrichment in 16:1 and a depletion of 18 carbon FAs (**Figure 3A, Supplemental Figure 4**). Unsupplemented IVFs contained significantly less 18:1, 18:2 versus P7 and P67 mouse fats, and slightly less 18:0 and 18:3 than P67 mouse (but similar levels to P7 mouse). 16:0 was observed at similar levels to P67 mouse, while being lower than P7 mouse. Lower carbon FAs were found at higher levels in unsupplemented IVFs (6:0, 14:1), but these FAs were also occasionally enriched in P7 mouse (14:0, 12:0, 10:0). The addition of lipids (0-1,000 μg/ml Intralipid) during cell culture altered cellular TAGs dose dependently, generally bringing FA compositions closer to that of the native fats with a reduction in 16:1. The increases within IVFs of 16:0 and 18 carbon FAs (18:0, 18:1, 18:2, 18:3) correlated with the composition of soybean oil, the major constituent of Intralipid (aside from water) (**Supplemental Table 3**)^50^. Principal component analysis (PCA) shows lipid supplementation modifying *in vitro* murine TAGs closer towards their native equivalents, particularly P7 mouse (**Supplemental Figure 5**). When looking at TAG FA saturation, samples did not vary greatly in terms of monounsaturated fatty acids (MUFAs), except for P7 mouse fat (**Figure 3C**). Unsupplemented IVFs contained low levels (2.9% ± 0.1) of polyunsaturated FAs (PUFAs), but this was increased to 12.3% (± 2.7) with 1000 μg/ml of Intralipid, similar to that of P7 mouse. PUFA increases for Intralipid IVFs were mirrored by a decrease in saturated FAs (SFAs), going from 48.2% (± 0.9, unsupplemented IVFs) to 39.9% (± 3.0, 1000 μg/ml Intralipid).

**Figure 4.**
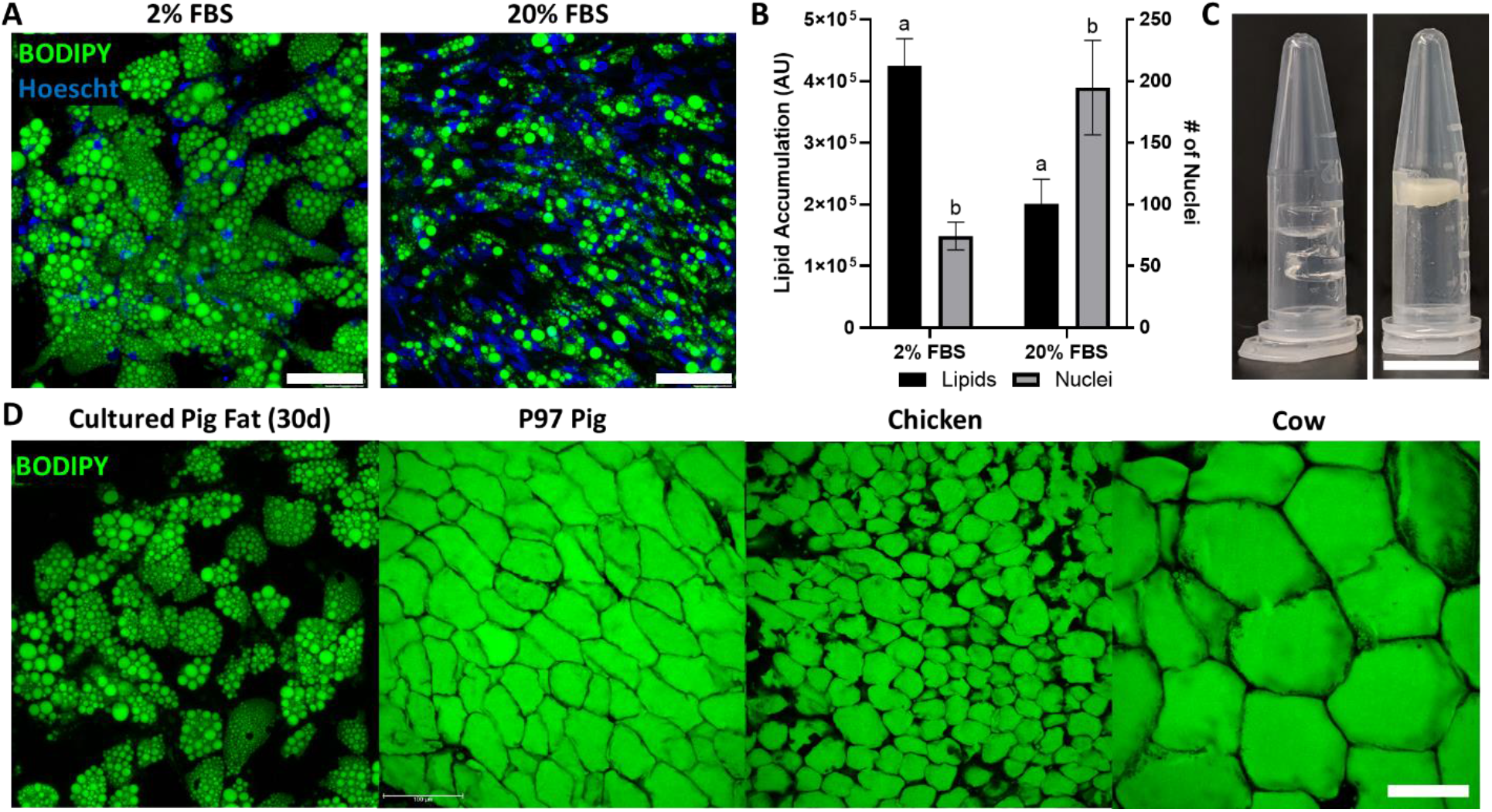
Porcine adipocytes grown to produce macroscale cultured fat. **(A)** Pig DFAT cells cultured under adipogenic conditions containing 2% or 20% FBS for 30 days. Fat cells are stained for lipid using BODIPY (green) and for cell nuclei using Hoescht 33342 (blue). Scale bars 100 μm. **(B)** Lipid and cell number quantification of 30-day pig adipocytes, based on BODIPY and Hoescht 33342 staining, respectively. AU stands for arbitrary units. Results tagged with ‘a’ represent a difference of p ≤ 0.0001, while ‘b’ represents p ≤ 0.01. **(C, Left)** Cell-free 1.6% alginate gels on the left. **(C, Right)** Porcine adipocytes (differentiated in 2% FBS media) mixed with alginate (1.6% final concentration) to form bulk cultured fat. Scale bar represents 1 mm. **(D)** Lipid staining images (BODIPY) of 30-day *in vitro* porcine adipocytes, juxtaposed with native adipocytes from a 97-day old (P97) pig, as well as a chicken and a cow. Scale bar 100 μm for all four images.

**Figure 5.**
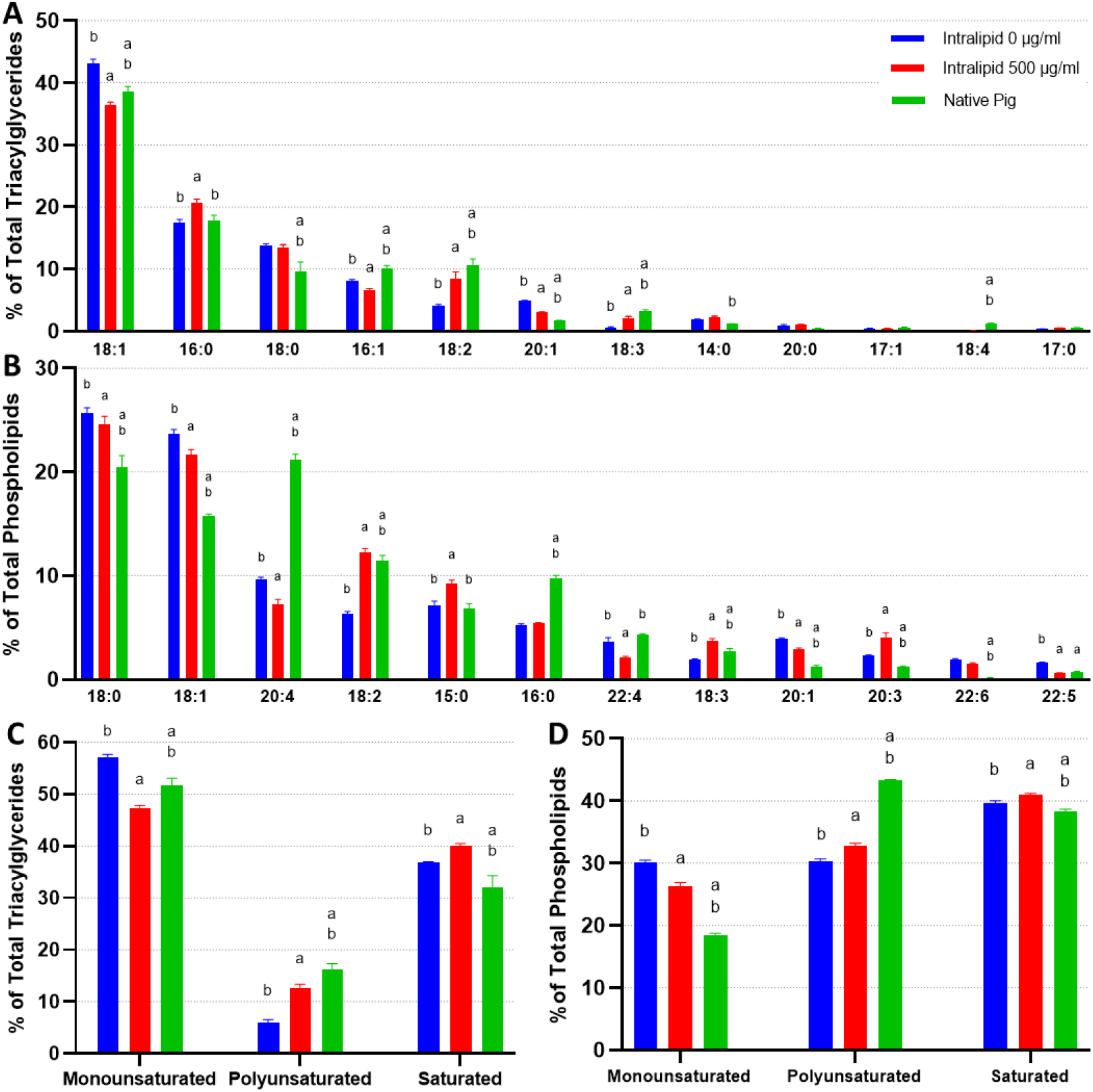
The fatty acid composition of **(A)** <u>triacylglycerides</u> and **(B)** phospholipids from *in vitro* (12 days of adipogenesis, 0-500 μg/ml Intralipid) and native <u>porcine</u> fats. The top 12 most prevalent fatty acids across all sample groups are shown. Tables containing the proportions of all detected fatty acids are available in **Supplemental Tables 6 and 7. (C)** and **(D)** represent fatty acids categorized by degree of saturation for triacylglycerides and phospholipids respectively. The top-down sample order in the legend is the same as the left-right order of the samples in the column charts. ‘a’ and ‘b’ represent a difference of p ≤ 0.05 versus 0 μg/ml Intralipid and 500 μg/ml Intralipid respectively. TAGs and phospholipids versus total lipids are available in **Supplemental Figure 10**.

In addition to TAGs, we also looked at phospholipid FA compositions due to their potential role in the flavor of meat^51,52^. Unsupplemented IVFs were similarly high in 16:1s as observed in the TAGs (**Figure 3B**). 16:0 levels were high and above that of P67 mouse, but similar to P7 mouse. 18:1 levels were lower than P67 mouse, but higher than P7 mouse. Unsupplemented IVFs contained noticeably fewer 18:0s and 20:4s than native fats. Lipid supplementation had a more muted effect on IVF phospholipids, with little to no changes in 16:0 contrary to the changes observed in the TAGs. Changes in MUFAs were observed, with 18:1 increasing 3.9% (± 0.6) and 16:1 decreasing 8.3% (± 1.4) from 0 to 1000 μg/ml Intralipid respectively. From looking at phospholipid FA saturation, the main changes from lipid supplementation are a decrease in MUFAs, with a corresponding increase in SFAs (**Figure 3D**). Overall though, lipid supplementation results in a decrease in IVF SFAs, as TAGs make up the majority of adipocyte lipids (**Supplemental Figure 6**).

### Porcine adipocytes can be grown *in vitro* and aggregated into 3D cultured fat

We additionally investigated whether it was possible to grow porcine adipocytes for aggregation into macroscale cultured fat. Primary porcine preadipocytes were observed to readily accumulate lipid (**Figure 4A**). As animal component-free culture media is key to cultivated meat, we also investigated reducing serum in our culture media. Interestingly, sample groups cultured in adipogenic media containing 2% FBS contained over 2X fewer cells, yet had over 2X the lipid accumulation, versus groups with 20% FBS (**Figure 4A, B**). Similar effects were seen in preliminary experiments with 0% FBS (**Supplemental Figure 7**). After 30 days of adipogenesis, cultured porcine adipocytes were also successfully combined with alginate to form macroscale cultured fat constructs (**Figure 4C**). When compared to native adipose from food animals (pig, chicken, cow), lipids within cultured porcine fats were multilocular, unlike native adipose from food animals (pig, chicken, cow) (**Figure 4D**).

### *In vitro* and native pig fat compositions exhibit similarities and can be tuned via lipid supplementation during cell culture

When exploring the effects of Intralipid on primary porcine adipocytes, we observed that lipid supplementation greatly increased their lipid accumulation, in contrast to 3T3-L1s (**Supplemental Figure 8**). Due to this, we also performed lipidomics to see if primary adipocytes also differed in terms of fat composition. Unlike 3T3-L1s, *in vitro* and native porcine fats shared many of their most abundant FAs, especially within their TAGs (**Figure 5A, B**). The only major difference observed was a lack of 20:4 (>10% difference) within the phospholipids, with smaller differences observed for 18:0, 18:1, 16:0 also within the phospholipids. Mild deficiencies of 18:2 and 18:3, as well as a slight abundance of 20:1, were observed across TAGs and phospholipids. Intralipid supplementation appeared to rescue 18:2 and 18:3 levels (while also decreasing 20:1), but it also caused the FA contents of *in vitro* porcine adipocytes to deviate in other cases (e.g., 15:0, 16:1, 20:3, 20:4, 22:4). This can be observed when looking at PCAs and FA saturation for TAGs and phospholipids, where Intralipid appears to both increase and decrease *in vitro* adipocyte similarity to native cells depending on the principal component (**Figure 5C, D, Supplemental Figure 9**).

## Discussion

Our goal was to generate macroscale fat tissue for food applications. To achieve this, we circumvented the limitations of macroscale tissue engineering and 3D cell culture by first culturing murine and porcine adipocytes on 2D surfaces with ample access to the culture media, then subsequently aggregating the cells into macroscale 3D tissues after *in vitro* adipogenesis. This approach is feasible for food applications, as opposed to regenerative medicine needs, because cell viability does not need to be maintained once the final macroscale fat tissue is produced.

We were first able to generate considerable amounts of highly lipid-laden murine adipocytes bearing morphological resemblance to native adipose tissues. Most studies that investigate 3T3-L1 adipogenesis in 2D are of limited culture duration (≤15 days), with lipid accumulation manifesting as small lipid droplets^53–59^. Here, we show that extended 2D culture (30 days) of *in vitro* adipocytes resulted in the emergence of larger lipid droplets, a key trait of white adipose tissue. Abundant lipids were also observed in porcine adipocytes, especially with 0-2% serum. However, lipid accumulation was multilocular. This morphology likely represents an immature white adipocyte phenotype, as Yorkshire pigs lack a functional UCP1 gene^60,61^. Optimizations to the *in vitro* environment and culture regime may help enhance lipid accumulation and promote a unilocular phenotype^62,63^. For example, in this study we found that Intralipid supplementation drastically increased porcine cell adipogenesis.

After *in vitro* adipocyte generation, mechanical aggregation of the cells during cell scraping immediately produced what looked like fat tissue. Murine adipocytes were then combined with food-relevant binders to maintain aggregation and tissue integrity. Uniaxial compression testing of alginate-based cultured fats revealed similar stress-strain behaviors to native fats, while mTG crosslinked adipocytes exhibited weaker resistance to compression closer to rendered fats. As mTG has been shown to greatly strengthen protein-rich materials, the weaker mTG-fats may be due to a lack of extracellular protein available for crosslinking amongst the aggregated *in vitro* adipocytes^64^. Thus, to strengthen mTG-fats it may be beneficial to mix in additional proteins such as soy^65^. It would also be interesting to investigate whether upregulating ECM deposition would improve mTG crosslinking, especially since ECM-promoting culture media components like ascorbic acid have also been shown to improve *in vitro* adipogenesis^66–70^. Ultimately, the mechanical properties of aggregated cultured fat tissues were tunable based on the binding technique used (alginate vs mTG), as well as via the concentration of binder (0.8 vs 1.6% alginate).

When investigating the TAG compositions of *in vitro-*grown fats, we found murine 3T3-L1 cells to differ from native fats, notably being deficient in 18:1 and 18:2 while being enriched in 16:1 and shorter chained SFAs. It has been shown that rodents lacking dietary oleic acid (18:1 ω-9) compensated with higher levels of 16:1, as endogenous production of oleic acid is insufficient to achieve the amounts typically found in normal tissue^71^. Primary porcine adipocytes on the other hand achieved a closer FA profile their native equivalents, though slightly lacking in certain FAs such as 18:2. The scarce presence of 18:2 for both murine and porcine cells is likely due an inability for mammals to synthesize linoleic acid (18:2 ω-6), which may highlight the importance of its supplementation during *in vitro* culture^72,73^. Indeed, Intralipid treatment (44-62% linoleic acid) enriched 18:2 and raised overall PUFA levels for all *in vitro* adipocytes, with murine cells additionally achieving more similarity to native mouse fats in terms of 16:1 and 18:1^50^. The increase in 18:3 from Intralipid across murine and porcine adipocytes represents a potential enrichment in omega 3 PUFAs, as 18:3 in soybean oil is alpha-linoleic acid. These changes suggest that lipid supplementation may be useful for tuning *in vitro* adipocyte TAGs to address deficiencies in particular FAs, enhance specific FAs, or more closely match the composition of native tissues.

As phospholipids have been reported to contribute meat flavor, we additionally looked at IVF phospholipid FAs^51,52^. *In vitro* murine and porcine adipocytes contained fewer 20:4 and 18:2 FAs relative to their native equivalents. As linoleic acid (18:2) is a precursor to arachidonic acid (20:4 ω-6), it was possible that low 20:4 was due to a lack of 18:2. However, Intralipid treated porcine IVFs did not show increased 20:4 despite having increases in 18:2. 20:4 levels in porcine IVFs were not uncommonly low though, achieving the same value as reported elsewhere for pig fat (1.4% of all FAs)^74^. High 20:4 in our native pig samples could be related to how native pork fat FAs are very pliable based on the animal’s diet^75^. Ultimately, optimal intracellular 20:4 levels are unclear, as arachidonic acid has been linked with both inflammation and enhanced meat flavor^76–78^. If desired, raising 20:4 may be possible through supplementation during cell culture, or via genetic interventions^78–80^. For murine IVF phospholipids, Intralipid did not raise 18:2 despite its high linoleic acid content^50^. This could be due to genetic influence over phospholipid FAs and our use of the 3T3-L1 cell line (less representative of native adipocytes). It has been reported that different strains of mice can have varying phospholipid compositions when fed the same diet^81^.

Generating cell-cultured animal fat via the aggregation of *in vitro* adipocytes unlocks a simplified approach of producing bulk cultured fat with lower costs and direct scalability, addressing a key obstacle in cultivated meat production and contributing to the paradigm of food production using cell culture
techniques. In addition to being cell-type and species agnostic as shown in this study, the adipocyte aggregation concept should be applicable to many cell cultivation techniques beyond laboratory-scale 2D culture. Scalable cell manufacturing techniques (e.g., hollow fiber, microcarrier-based stirred suspension bioreactors) could grow large amounts of adipocytes to be combined at the end of culture to form macroscale fat after adipocytes have completed lipid accumulation and no longer require continued cell culture^82,83^. Directly culturing adipocytes in suspension may even be possible, as demonstrated with 3T3-L1s^84^. These approaches offer potential paths to the overall production of meat without animal slaughter, while potentially being more sustainable^85–89^. Incorporating cultured fat into plant-based meats also permits the creation of hybrid products, where sustainable and low-cost plant ingredients are enhanced by complex species-specific flavors from added animal fat cells.

## Acknowledgements

We thank ARPA-E (DE-AR0001233), the NIH (P41EB027062), New Harvest and the United States Department of Defense (DoD) through the National Defense Science & Engineering Graduate (NDSEG) Fellowship Program. We are also grateful to Dr. John Asara and the Mass Spectrometry Core Facility at the Beth Israel Deaconess Medical Center for assistance with lipidomics, Sawnaz Shaidani for assistance with BioRender, and Kevin Zhang for help with figures. Our graphical abstract was created with BioRender.com.

## Supplemental Tables

**Supplemental Table 1.** Proportion of nuclei positive for the adipogenic transcription factor PPARγ in 3T3-L1 adipocytes grown under adipogenic conditions for 15 days.

| PPARγ Positive Nuclei (%) | |
| --- | --- |
| Mean | 82.53 |
| Standard Deviation | 5.18 |

**Supplemental Table 2.** A frequency distribution table of 15- and 30-day cultured adipocyte lipid droplet diameters compared using a chi-square test (degrees of freedom = 8).

| Lipid Droplet Diameters (%) |  |  |
| --- | --- | --- |
| Diameter (μm) | 15 Days | 30 Days |
| 0-5 | 14.94 | 16.78 |
| 5-10 | 65.78 | 45.67 |
| 10-15 | 12.68 | 16.38 |
| 15-20 | 3.96 | 8.57 |
| 20-25 | 1.59 | 4.77 |
| 25-30 | 0.74 | 3.65 |
| 30-35 | 0.24 | 1.79 |
| 35-40 | 0.06 | 1.28 |
| $\geq 40$ | 0.01 | 1.12 |
| Chi-Square Value: 1052, p = <0.0001 |  |  |

**Supplemental Table 3.**
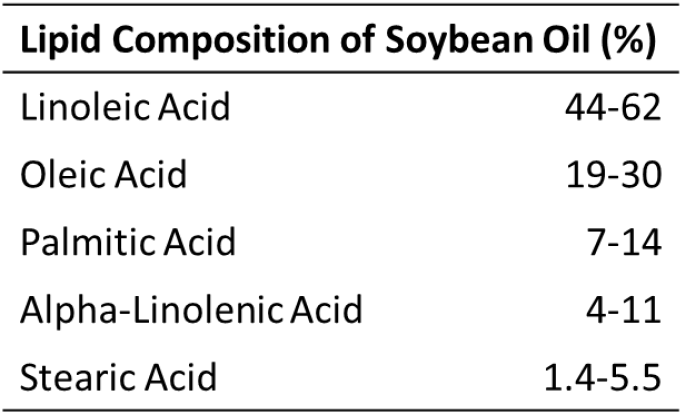
The major fatty acids present in soybean oil (the major component of Intralipid, other than water), according to the package insert supplied with Intralipid 20% (Intravenous Fat Emulsion).

| Lipid Composition of Soybean Oil (%) |  |
| --- | --- |
| Linoleic Acid | 44-62 |
| Oleic Acid | 19-30 |
| Palmitic Acid | 7-14 |
| Alpha-Linolenic Acid | 4-11 |
| Stearic Acid | 1.4-5.5 |

**Supplemental Table 4.** The full fatty acid compositions (mean % of total of <u>triacylglycerides</u> measured, n=3) of *in vitro*-(30 days of adipogenesis) and *in vivo*-grown <u>murine</u> fats.

|  | Intra-0 | Intra-500 | Intra-1000 | P7 Mouse | P67 Mouse | 21:0 | 0.021% | 0.037% | 0.085% | 0.013% | 0.000% |
| --- | --- | --- | --- | --- | --- | --- | --- | --- | --- | --- | --- |
| 16:1 | 32.751% | 28.668% | 19.831% | 17.439% | 15.818% | 25:1 | 0.060% | 0.001% | 0.005% | 0.002% | 0.087% |
| 16:0 | 17.113% | 19.644% | 21.906% | 24.107% | 16.299% | 26:1 | 0.040% | 0.002% | 0.022% | 0.065% | 0.018% |
| 18:1 | 6.138% | 13.699% | 24.548% | 20.239% | 29.174% | 26:0 | 0.001% | 0.003% | 0.022% | 0.063% | 0.052% |
| 18:2 | 2.496% | 5.339% | 7.449% | 7.619% | 16.991% | 23:0 | 0.004% | 0.018% | 0.089% | 0.010% | 0.000% |
| 14:0 | 6.844% | 5.640% | 2.824% | 5.658% | 0.534% | 20:5 | 0.000% | 0.024% | 0.072% | 0.019% | 0.000% |
| 6:0 | 6.884% | 4.437% | 2.215% | 1.755% | 0.196% | 25:0 | 0.007% | 0.003% | 0.038% | 0.021% | 0.040% |
| 12:0 | 3.718% | 1.036% | 1.084% | 6.757% | 0.782% | 14:4 | 0.003% | 0.015% | 0.088% | 0.000% | 0.000% |
| 18:0 | 0.526% | 1.198% | 3.900% | 1.429% | 4.780% | 22:4 | 0.001% | 0.011% | 0.042% | 0.019% | 0.027% |
| 15:0 | 3.437% | 4.083% | 2.267% | 0.525% | 0.915% | 23:1 | 0.000% | 0.016% | 0.063% | 0.000% | 0.021% |
| 14:1 | 5.585% | 2.153% | 0.788% | 1.504% | 1.191% | 21:1 | 0.002% | 0.006% | 0.054% | 0.003% | 0.031% |
| 10:0 | 3.814% | 0.727% | 0.389% | 4.587% | 0.358% | 30:0 | 0.001% | 0.065% | 0.026% | 0.000% | 0.000% |
| 4:0 | 3.216% | 2.489% | 1.417% | 1.217% | 1.393% | 22:2 | 0.004% | 0.045% | 0.034% | 0.004% | 0.000% |
| 18:3 | 0.214% | 0.997% | 2.923% | 0.500% | 2.984% | 22:5 | 0.000% | 0.015% | 0.036% | 0.008% | 0.015% |
| 17:1 | 2.495% | 2.069% | 1.028% | 0.603% | 1.095% | 10:4 | 0.003% | 0.019% | 0.043% | 0.006% | 0.000% |
| 10:1 | 0.837% | 1.077% | 0.516% | 0.863% | 0.413% | 30:1 | 0.002% | 0.036% | 0.026% | 0.000% | 0.001% |
| 17:0 | 0.801% | 0.715% | 1.067% | 0.491% | 0.394% | 15:1 | 0.029% | 0.017% | 0.008% | 0.007% | 0.000% |
| 8:0 | 0.351% | 1.074% | 0.490% | 0.798% | 0.118% | 14:2 | 0.000% | 0.018% | 0.007% | 0.000% | 0.012% |
| 9:0 | 0.821% | 0.918% | 0.848% | 0.091% | 0.050% | 10:3 | 0.003% | 0.007% | 0.016% | 0.006% | 0.000% |
| 24:1 | 0.030% | 0.055% | 0.074% | 0.201% | 1.814% | 12:4 | 0.000% | 0.005% | 0.022% | 0.000% | 0.000% |
| 12:1 | 0.381% | 0.188% | 0.355% | 1.098% | 0.072% | 29:0 | 0.000% | 0.010% | 0.001% | 0.000% | 0.014% |
| 11:0 | 0.319% | 0.892% | 0.190% | 0.137% | 0.065% | 7:0 | 0.000% | 0.007% | 0.013% | 0.000% | 0.000% |
| 19:0 | 0.146% | 0.274% | 0.229% | 0.226% | 0.543% | 12:3 | 0.000% | 0.003% | 0.017% | 0.000% | 0.000% |
| 13:0 | 0.022% | 0.412% | 0.192% | 0.129% | 0.540% | 8:1 | 0.001% | 0.001% | 0.012% | 0.001% | 0.000% |
| 24:0 | 0.011% | 0.046% | 0.140% | 0.084% | 0.921% | 5:0 | 0.007% | 0.001% | 0.002% | 0.002% | 0.000% |
| 20:1 | 0.078% | 0.280% | 0.123% | 0.312% | 0.389% | 22:3 | 0.000% | 0.001% | 0.006% | 0.001% | 0.000% |
| 9:2 | 0.003% | 0.337% | 0.688% | 0.008% | 0.000% | 11:2 | 0.000% | 0.000% | 0.000% | 0.002% | 0.005% |
| 11:1 | 0.236% | 0.432% | 0.138% | 0.114% | 0.079% | 6:1 | 0.001% | 0.000% | 0.006% | 0.000% | 0.000% |
| 22:0 | 0.079% | 0.305% | 0.314% | 0.213% | 0.071% | 11:3 | 0.000% | 0.001% | 0.006% | 0.000% | 0.000% |
| 20:4 | 0.028% | 0.035% | 0.190% | 0.461% | 0.194% | 13:2 | 0.002% | 0.001% | 0.001% | 0.002% | 0.000% |
| 22:1 | 0.231% | 0.007% | 0.152% | 0.106% | 0.348% | 28:1 | 0.000% | 0.000% | 0.000% | 0.000% | 0.005% |
| 22:6 | 0.003% | 0.068% | 0.053% | 0.052% | 0.304% | 27:0 | 0.000% | 0.000% | 0.000% | 0.000% | 0.004% |
| 19:1 | 0.029% | 0.130% | 0.124% | 0.022% | 0.152% | 18:5 | 0.000% | 0.000% | 0.004% | 0.000% | 0.000% |
| 20:3 | 0.001% | 0.007% | 0.014% | 0.137% | 0.287% | 27:1 | 0.000% | 0.000% | 0.002% | 0.000% | 0.000% |
| 12:2 | 0.060% | 0.053% | 0.319% | 0.002% | 0.009% | 14:3 | 0.000% | 0.000% | 0.000% | 0.000% | 0.002% |
| 20:2 | 0.002% | 0.035% | 0.101% | 0.119% | 0.140% | 16:2 | 0.000% | 0.001% | 0.001% | 0.000% | 0.000% |
| 20:0 | 0.080% | 0.026% | 0.128% | 0.038% | 0.071% | 20:6 | 0.000% | 0.000% | 0.000% | 0.001% | 0.000% |
| 10:2 | 0.004% | 0.049% | 0.041% | 0.035% | 0.049% | 29:1 | 0.000% | 0.000% | 0.000% | 0.000% | 0.001% |
| 18:4 | 0.005% | 0.005% | 0.016% | 0.001% | 0.133% | 11:4 | 0.000% | 0.000% | 0.000% | 0.000% | 0.000% |
| 24:2 | 0.018% | 0.011% | 0.062% | 0.067% | 0.001% | 16:4 | 0.000% | 0.000% | 0.000% | 0.000% | 0.000% |
| 21:0 | 0.021% | 0.037% | 0.085% | 0.013% | 0.000% | 28:0 | 0.000% | 0.000% | 0.000% | 0.000% | 0.000% |

**Supplemental Table 5.** The full fatty acid compositions (mean % of total of <u>phospholipids</u> measured, n=3) of *in vitro*-(30 days of adipogenesis) and *in vivo*-grown <u>murine</u> fats.

|  | Intra-0 | Intra-500 | Intra-1000 | P7 Mouse | P67 Mouse | 19:1 | 0.000% | 0.000% | 0.000% | 0.000% | 0.083% |
| --- | --- | --- | --- | --- | --- | --- | --- | --- | --- | --- | --- |
| 16:0 | 27.884% | 29.787% | 29.602% | 28.741% | 13.725% | 27:0 | 0.000% | 0.012% | 0.051% | 0.000% | 0.000% |
| 18:1 | 17.907% | 21.949% | 22.153% | 13.207% | 28.005% | 19:0 | 0.003% | 0.012% | 0.015% | 0.002% | 0.025% |
| 16:1 | 28.914% | 20.606% | 24.173% | 1.426% | 9.952% | 26:1 | 0.000% | 0.016% | 0.005% | 0.000% | 0.000% |
| 18:0 | 4.793% | 8.729% | 6.104% | 16.964% | 14.117% | 33:1 | 0.000% | 0.000% | 0.000% | 0.000% | 0.019% |
| 20:4 | 5.201% | 4.475% | 4.450% | 22.979% | 9.744% | 25:0 | 0.000% | 0.000% | 0.000% | 0.000% | 0.015% |
| 18:2 | 2.417% | 1.397% | 1.586% | 4.496% | 5.711% | 14:3 | 0.000% | 0.003% | 0.011% | 0.000% | 0.000% |
| 22:6 | 2.280% | 1.593% | 1.739% | 3.352% | 2.794% | 28:0 | 0.000% | 0.002% | 0.002% | 0.000% | 0.004% |
| 17:1 | 5.595% | 0.356% | 0.725% | 0.061% | 3.099% | 12:0 | 0.000% | 0.000% | 0.000% | 0.000% | 0.001% |
| 15:0 | 1.743% | 1.841% | 1.414% | 2.873% | 0.906% | 23:0 | 0.000% | 0.000% | 0.000% | 0.000% | 0.001% |
| 13:0 | 0.455% | 1.536% | 2.047% | 1.225% | 1.672% | 9:0 | 0.000% | 0.000% | 0.000% | 0.000% | 0.000% |
| 20:3 | 0.308% | 2.540% | 1.577% | 1.455% | 0.922% | 11:0 | 0.000% | 0.000% | 0.000% | 0.000% | 0.000% |
| 22:5 | 0.332% | 1.887% | 1.163% | 1.427% | 1.308% | 10:0 | 0.000% | 0.000% | 0.000% | 0.000% | 0.000% |
| 17:0 | 0.197% | 0.363% | 0.631% | 0.050% | 3.164% | 11:2 | 0.000% | 0.000% | 0.000% | 0.000% | 0.000% |
| 22:4 | 0.294% | 0.411% | 0.270% | 0.921% | 0.978% | 11:3 | 0.000% | 0.000% | 0.000% | 0.000% | 0.000% |
| 14:0 | 0.876% | 0.312% | 0.506% | 0.108% | 0.059% | 11:4 | 0.000% | 0.000% | 0.000% | 0.000% | 0.000% |
| 14:1 | 0.308% | 0.620% | 0.771% | 0.008% | 0.011% | 12:4 | 0.000% | 0.000% | 0.000% | 0.000% | 0.000% |
| 20:5 | 0.236% | 0.339% | 0.339% | 0.112% | 0.618% | 14:2 | 0.000% | 0.000% | 0.000% | 0.000% | 0.000% |
| 22:0 | 0.001% | 0.033% | 0.018% | 0.004% | 0.899% | 14:4 | 0.000% | 0.000% | 0.000% | 0.000% | 0.000% |
| 18:3 | 0.081% | 0.077% | 0.131% | 0.495% | 0.149% | 15:1 | 0.000% | 0.000% | 0.000% | 0.000% | 0.000% |
| 24:0 | 0.012% | 0.109% | 0.056% | 0.010% | 0.611% | 21:1 | 0.000% | 0.000% | 0.000% | 0.000% | 0.000% |
| 20:0 | 0.019% | 0.136% | 0.057% | 0.001% | 0.544% | 29:1 | 0.000% | 0.000% | 0.000% | 0.000% | 0.000% |
| 22:2 | 0.001% | 0.010% | 0.002% | 0.000% | 0.540% | 30:1 | 0.000% | 0.000% | 0.000% | 0.000% | 0.000% |
| 20:1 | 0.001% | 0.338% | 0.090% | 0.001% | 0.024% | 31:1 | 0.000% | 0.000% | 0.000% | 0.000% | 0.000% |
| 22:3 | 0.010% | 0.167% | 0.082% | 0.018% | 0.040% | 33:0 | 0.000% | 0.000% | 0.000% | 0.000% | 0.000% |
| 22:1 | 0.010% | 0.061% | 0.016% | 0.001% | 0.121% | 34:1 | 0.000% | 0.000% | 0.000% | 0.000% | 0.000% |
| 12:1 | 0.098% | 0.020% | 0.063% | 0.009% | 0.000% | 35:2 | 0.000% | 0.000% | 0.000% | 0.000% | 0.000% |
| 8:0 | 0.012% | 0.110% | 0.054% | 0.010% | 0.000% | 38:2 | 0.000% | 0.000% | 0.000% | 0.000% | 0.000% |
| 24:1 | 0.001% | 0.083% | 0.037% | 0.000% | 0.062% | 40:1 | 0.000% | 0.000% | 0.000% | 0.000% | 0.000% |
| 20:2 | 0.000% | 0.048% | 0.038% | 0.000% | 0.050% | 42:3 | 0.000% | 0.000% | 0.000% | 0.000% | 0.000% |
| 18:4 | 0.013% | 0.021% | 0.020% | 0.043% | 0.029% | 6:0 | 0.000% | 0.000% | 0.000% | 0.000% | 0.000% |

**Supplemental Table 6.** The full fatty acid compositions (mean % of total of <u>triacylglycerides</u> measured, n=3) of *in vitro*-(30 days of adipogenesis) and *in vivo*-grown <u>porcine</u> fats.

|  | Native Pig | 0 µg/ml Intralipid | 500 µg/ml Intralipid |  |  |  |  |
| --- | --- | --- | --- | --- | --- | --- | --- |
| 18:1 | 38.63% | 43.14% | 36.45% | 20:2 | 0.02% | 0.02% | 0.09% |
| 16:0 | 17.75% | 17.50% | 20.61% | 21:1 | 0.04% | 0.05% | 0.03% |
| 18:0 | 9.66% | 13.86% | 13.46% | 26:1 | 0.00% | 0.05% | 0.05% |
| 16:1 | 10.08% | 8.14% | 6.68% | 12:1 | 0.02% | 0.01% | 0.05% |
| 18:2 | 10.61% | 4.03% | 8.50% | 23:1 | 0.01% | 0.05% | 0.02% |
| 20:1 | 1.72% | 4.94% | 3.06% | 4:0 | 0.03% | 0.02% | 0.02% |
| 18:3 | 3.32% | 0.58% | 2.12% | 11:0 | 0.02% | 0.02% | 0.02% |
| 14:0 | 1.18% | 1.84% | 2.19% | 12:4 | 0.01% | 0.00% | 0.04% |
| 20:0 | 0.47% | 0.97% | 1.13% | 25:1 | 0.01% | 0.00% | 0.04% |
| 17:1 | 0.62% | 0.47% | 0.49% | 13:0 | 0.01% | 0.01% | 0.03% |
| 18:4 | 1.26% | 0.07% | 0.16% | 27:1 | 0.01% | 0.03% | 0.00% |
| 17:0 | 0.53% | 0.38% | 0.55% | 25:0 | 0.01% | 0.01% | 0.02% |
| 24:0 | 0.32% | 0.60% | 0.48% | 26:0 | 0.00% | 0.01% | 0.02% |
| 10:0 | 0.75% | 0.35% | 0.17% | 5:0 | 0.00% | 0.01% | 0.01% |
| 20:3 | 0.39% | 0.41% | 0.27% | 22:1 | 0.00% | 0.01% | 0.00% |
| 20:4 | 0.13% | 0.16% | 0.56% | 11:3 | 0.00% | 0.00% | 0.01% |
| 19:0 | 0.27% | 0.25% | 0.25% | 12:2 | 0.00% | 0.00% | 0.01% |
| 9:2 | 0.11% | 0.05% | 0.51% | 11:2 | 0.00% | 0.00% | 0.01% |
| 19:1 | 0.22% | 0.18% | 0.23% | 11:1 | 0.00% | 0.00% | 0.01% |
| 22:0 | 0.05% | 0.28% | 0.20% | 14:1 | 0.00% | 0.00% | 0.00% |
| 23:0 | 0.22% | 0.18% | 0.13% | 30:0 | 0.00% | 0.00% | 0.01% |
| 9:0 | 0.14% | 0.14% | 0.24% | 22:3 | 0.00% | 0.00% | 0.00% |
| 21:0 | 0.12% | 0.12% | 0.22% | 7:0 | 0.00% | 0.00% | 0.00% |
| 15:0 | 0.15% | 0.06% | 0.20% | 6:1 | 0.00% | 0.00% | 0.00% |
| 10:1 | 0.30% | 0.05% | 0.03% | 10:2 | 0.00% | 0.00% | 0.00% |
| 20:5 | 0.10% | 0.11% | 0.12% | 12:3 | 0.00% | 0.00% | 0.00% |
| 6:0 | 0.14% | 0.11% | 0.05% | 28:0 | 0.00% | 0.00% | 0.00% |
| 22:4 | 0.11% | 0.13% | 0.06% | 22:2 | 0.00% | 0.00% | 0.00% |
| 12:0 | 0.18% | 0.01% | 0.09% | 8:1 | 0.00% | 0.00% | 0.00% |
| 10:4 | 0.04% | 0.16% | 0.06% | 14:2 | 0.00% | 0.00% | 0.00% |
| 22:5 | 0.07% | 0.12% | 0.03% | 11:4 | 0.00% | 0.00% | 0.00% |
| 24:1 | 0.01% | 0.08% | 0.10% | 24:2 | 0.00% | 0.00% | 0.00% |
| 8:0 | 0.09% | 0.05% | 0.04% | 16:2 | 0.00% | 0.00% | 0.00% |
| 10:3 | 0.05% | 0.09% | 0.03% | 14:4 | 0.00% | 0.00% | 0.00% |
| 22:6 | 0.01% | 0.08% | 0.05% | 30:1 | 0.00% | 0.00% | 0.00% |

**Supplemental Table 7.** The full fatty acid compositions (mean % of total of <u>phospholipids</u> measured, n=3) of *in vitro*-(30 days of adipogenesis) and *in vivo*-grown <u>porcine</u> fats.

|  | Native Pig | 0 µg/ml Intralipid | 500 µg/ml Intralipid |  |  |  |  |
| --- | --- | --- | --- | --- | --- | --- | --- |
| **18:0** | 20.44% | 25.73% | 24.59% | **22:2** | 0.01% | 0.47% | 0.03% |
| **18:1** | 15.74% | 23.70% | 21.65% | **19:1** | 0.01% | 0.20% | 0.27% |
| **20:4** | 21.21% | 9.69% | 7.28% | **22:0** | 0.06% | 0.29% | 0.12% |
| **18:2** | 11.46% | 6.37% | 12.24% | **11:0** | 0.00% | 0.11% | 0.27% |
| **15:0** | 6.84% | 7.19% | 9.28% | **24:3** | 0.17% | 0.05% | 0.05% |
| **16:0** | 9.72% | 5.22% | 5.41% | **20:0** | 0.00% | 0.20% | 0.06% |
| **22:4** | 4.33% | 3.69% | 2.16% | **22:1** | 0.02% | 0.23% | 0.01% |
| **18:3** | 2.70% | 1.91% | 3.75% | **17:0** | 0.05% | 0.07% | 0.10% |
| **20:1** | 1.24% | 3.99% | 2.93% | **24:2** | 0.01% | 0.12% | 0.01% |
| **20:3** | 1.21% | 2.36% | 4.06% | **24:0** | 0.01% | 0.05% | 0.04% |
| **22:6** | 0.17% | 1.92% | 1.52% | **21:1** | 0.00% | 0.04% | 0.00% |
| **22:5** | 0.72% | 1.63% | 0.65% | **24:1** | 0.00% | 0.04% | 0.00% |
| **13:0** | 1.09% | 0.79% | 1.04% | **19:0** | 0.01% | 0.02% | 0.01% |
| **20:5** | 1.25% | 0.66% | 0.25% | **14:0** | 0.00% | 0.00% | 0.02% |
| **16:1** | 0.38% | 0.97% | 0.79% | **14:2** | 0.02% | 0.00% | 0.00% |
| **20:2** | 0.10% | 0.77% | 0.62% | **6:0** | 0.02% | 0.00% | 0.00% |
| **14:1** | 0.74% | 0.26% | 0.31% | **28:0** | 0.00% | 0.01% | 0.01% |
| **17:1** | 0.23% | 0.63% | 0.26% | **12:0** | 0.00% | 0.00% | 0.00% |
| **22:3** | 0.01% | 0.61% | 0.20% | **12:1** | 0.00% | 0.00% | 0.00% |

## Supplemental Figures

**Supplemental Figure 1.**
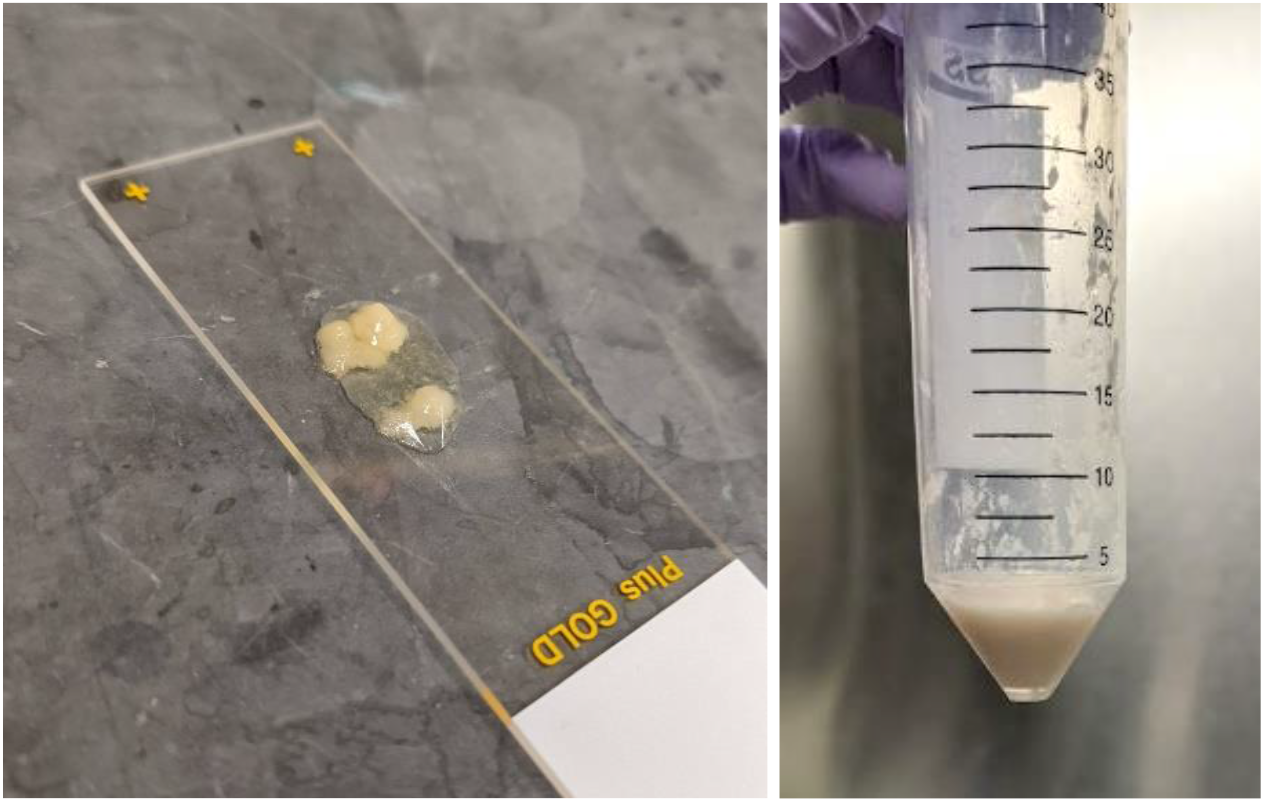
**(Left)** Cell-scraped adipocytes prepared on a glass slide (25×75 mm) with mounting media for subsequent microscopy. **(Right)** Cell scraped adipocytes from three T175 cell culture flasks collected in a 50 ml conical tube. The bulk aggregated cells take on an appearance similar to lipoaspirate.

**Supplemental Figure 2.**
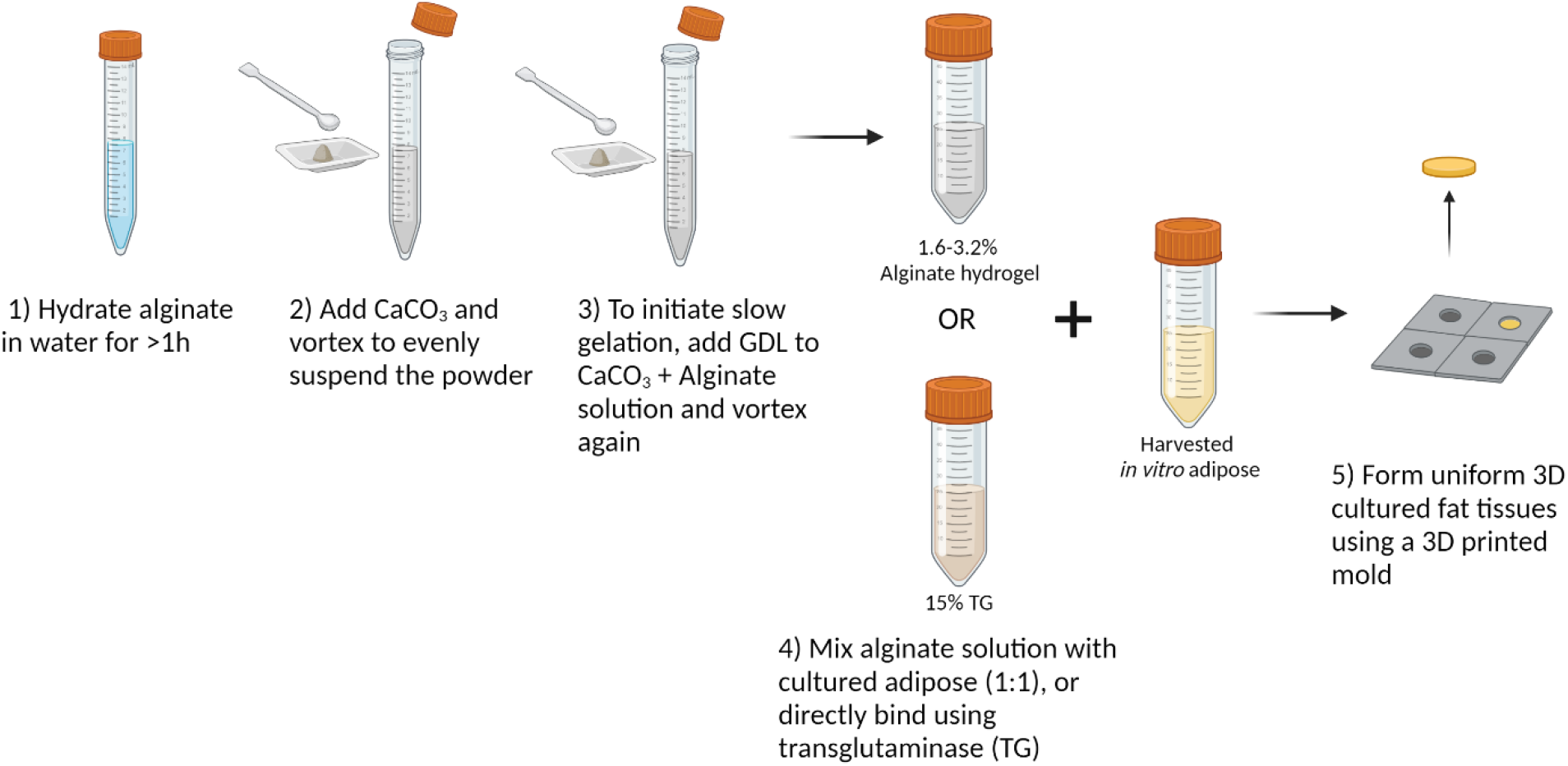
Detailed steps for producing macroscale 3D cultured fat constructs from aggregated adipocytes, using slow-gelling alginate or microbial transglutaminase as binders. Created using BioRender.com.

**Supplemental Figure 3.**
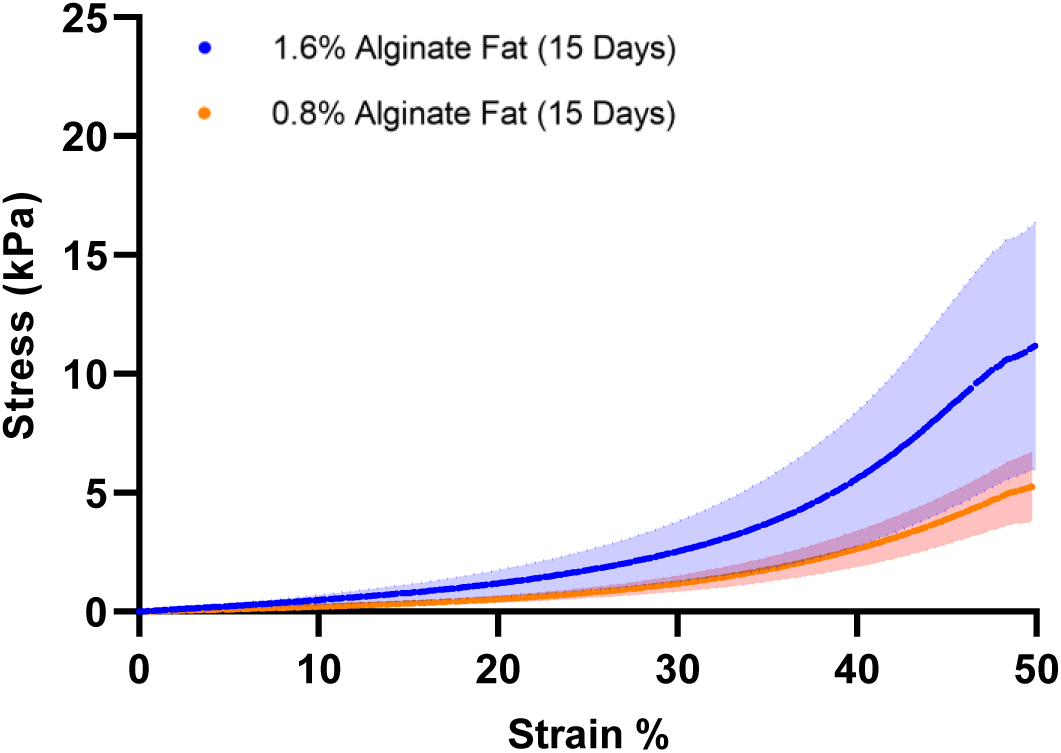
Uniaxial compression testing of macroscale cultured fat tissues produced from 0.8% and 1.6% (final concentration) alginate containing aggregated lipid-laden *in vitro* mouse adipocytes (15 days of adipogenesis).

**Supplemental Figure 4.**
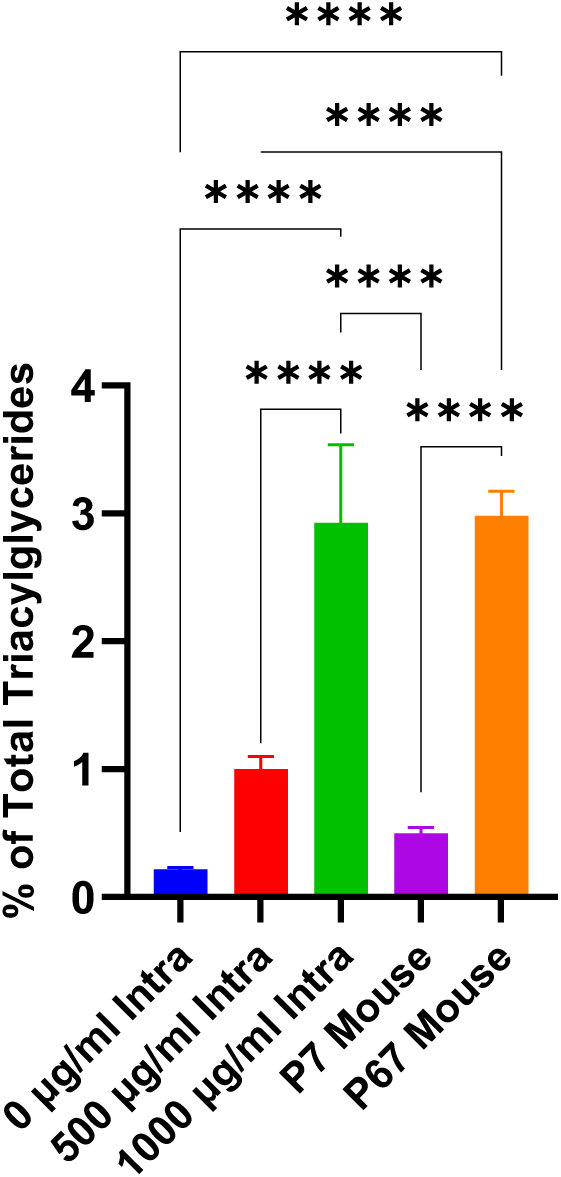
The 18:3 (triacylglyceride fraction) content of 3T3-L1 adipocytes adipogenically cultured for 30 days with various levels of fatty acid (Intralipid) supplementation, versus fat from 7-day and 67-day old mice. (****) denotes a p ≤0.0001 difference.

**Supplemental Figure 5.**
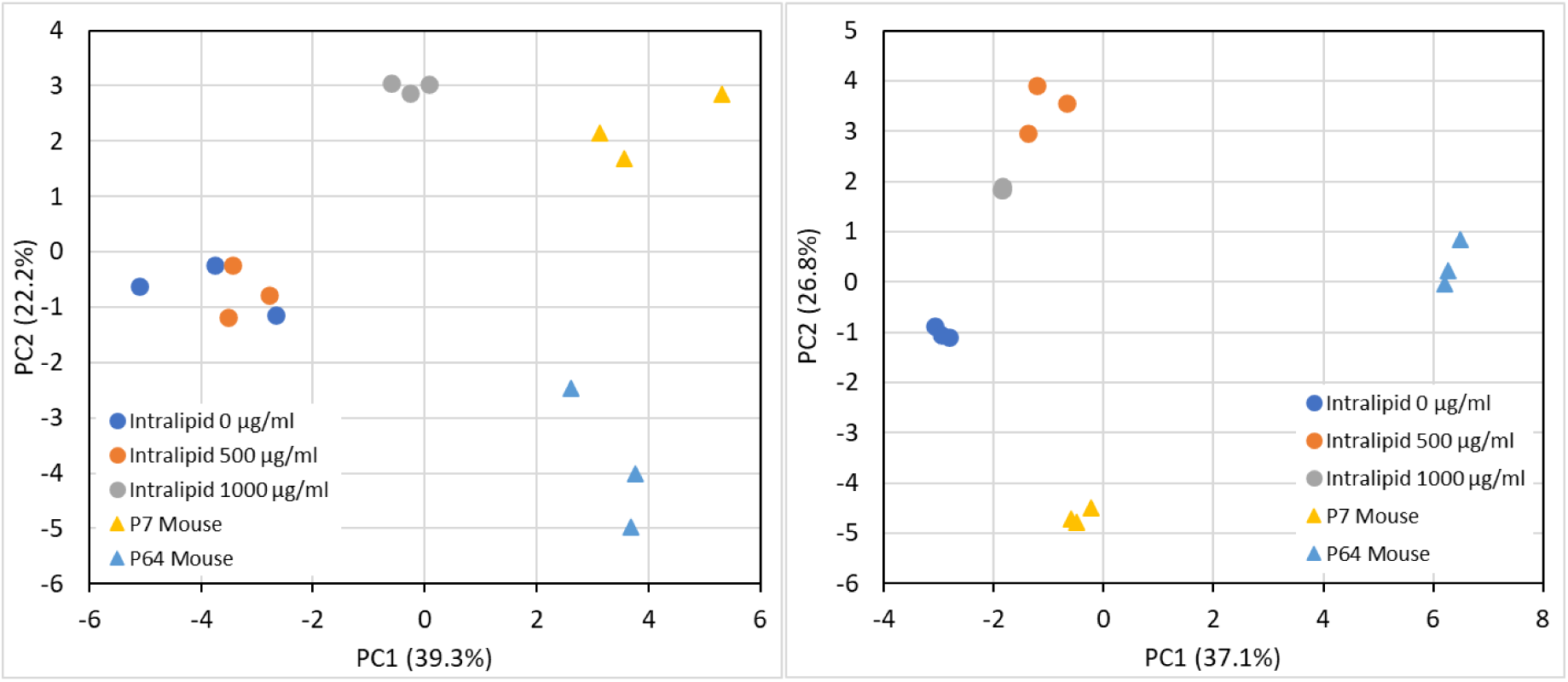
Principal component analyses of **(Left)** <u>triacylglyceride</u> (TAG) and **(Right)** phospholipid fatty acid compositions for *in vitro* and native <u>murine</u> fats (top 30 fatty acids). Circles (•) represent *in vitro* mouse fats while triangles (▴) represent native mouse fats.

**Supplemental Figure 6.**
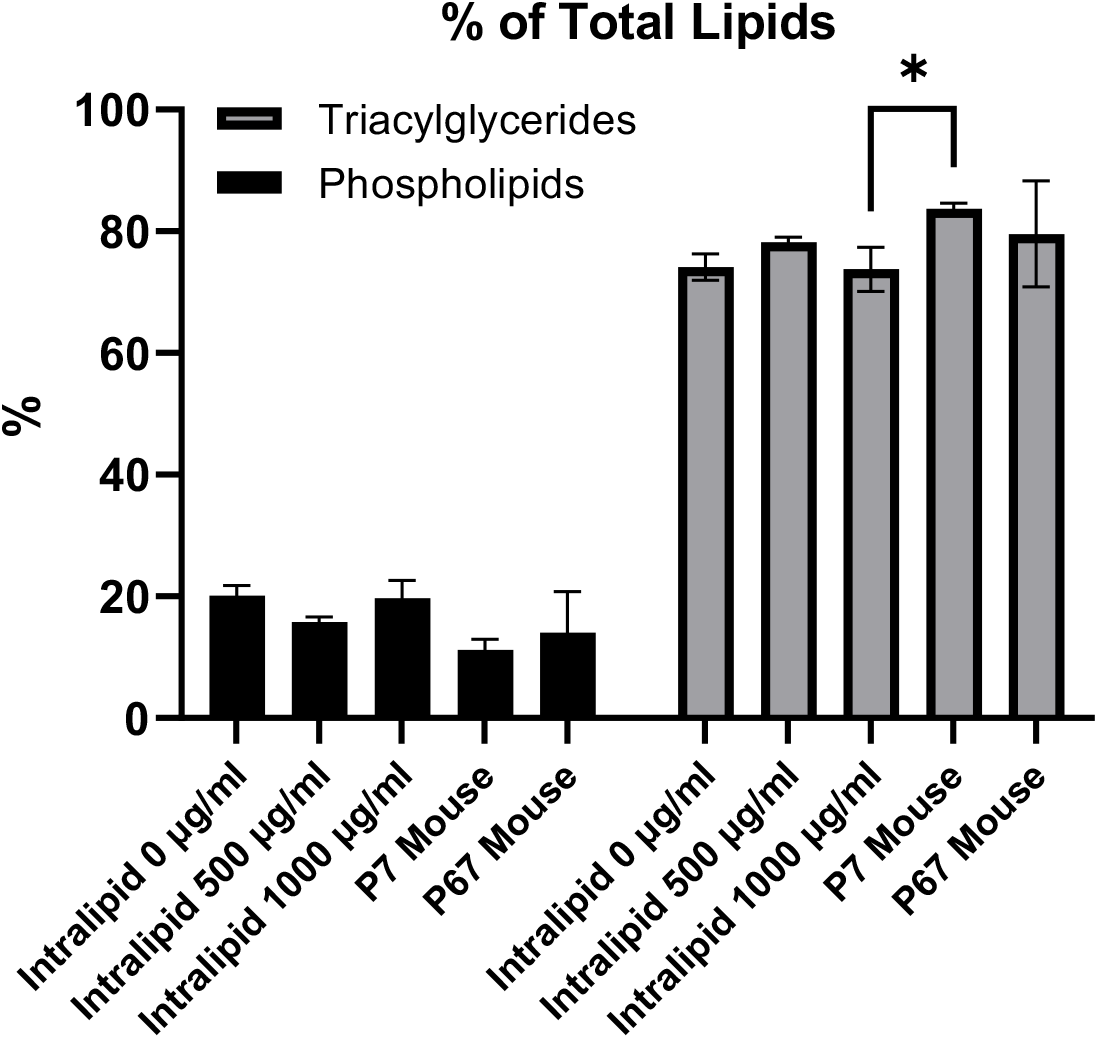
The proportion of triacylglycerides and phospholipids as a percentage of total intracellular lipids, for *in vitro* and native <u>murine</u> fats. (*) represents p < 0.05.

**Supplemental Figure 7.**
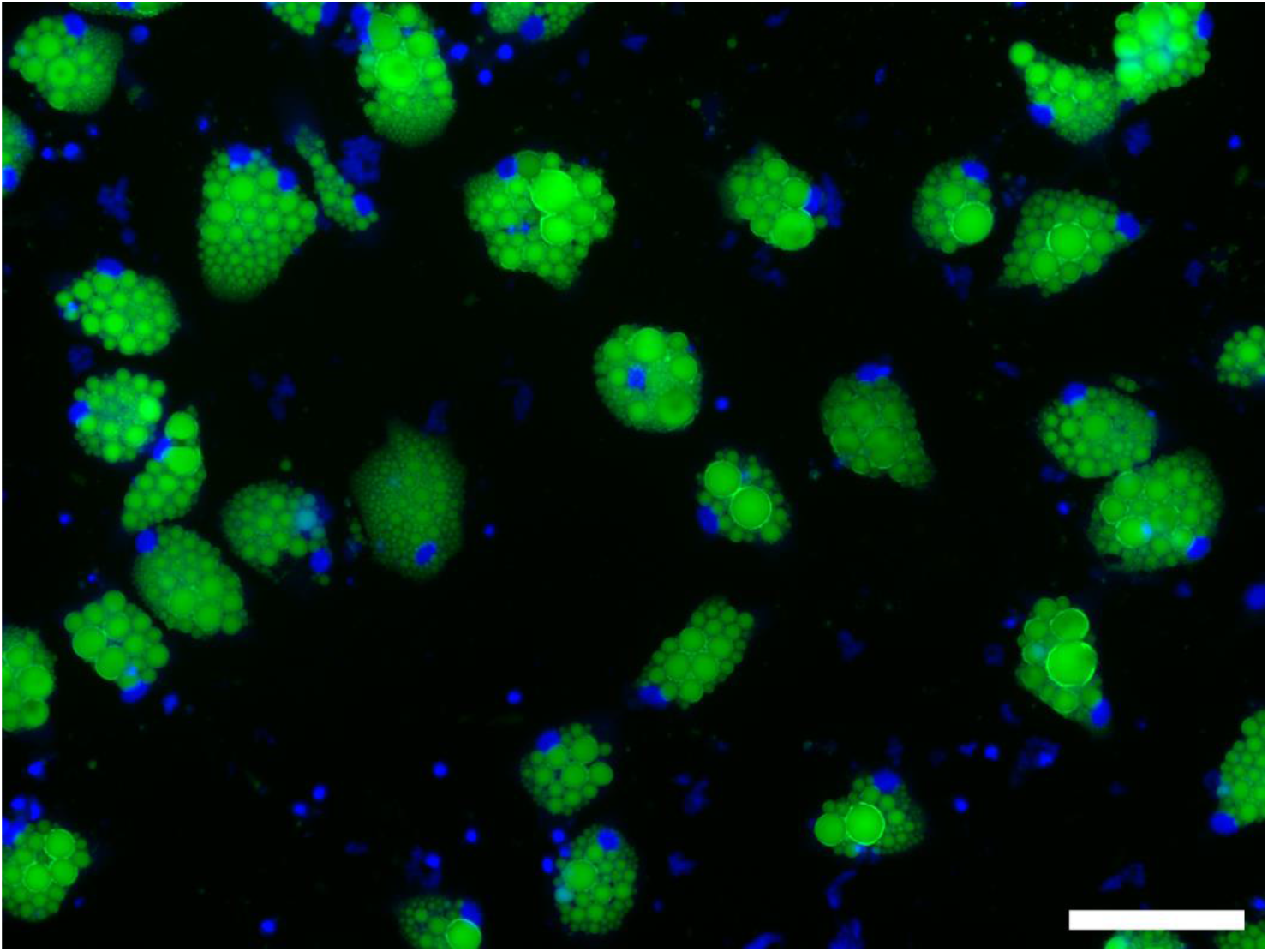
Pig DFAT cells cultured under adipogenic conditions with 0% FBS containing media for 30 days. Cells were grown similarly to adipocytes cultured in adipogenic media containing 2% FBS, except the media did not contain GlutaMAX or FBS. The induction and lipid accumulation phases lasted 6 days and 24 days respectively. Lipid staining is BODIPY (green) and nuclei staining is Hoescht 33342 (blue). Scale bar represents 100 μm.

**Supplemental Figure 8.**
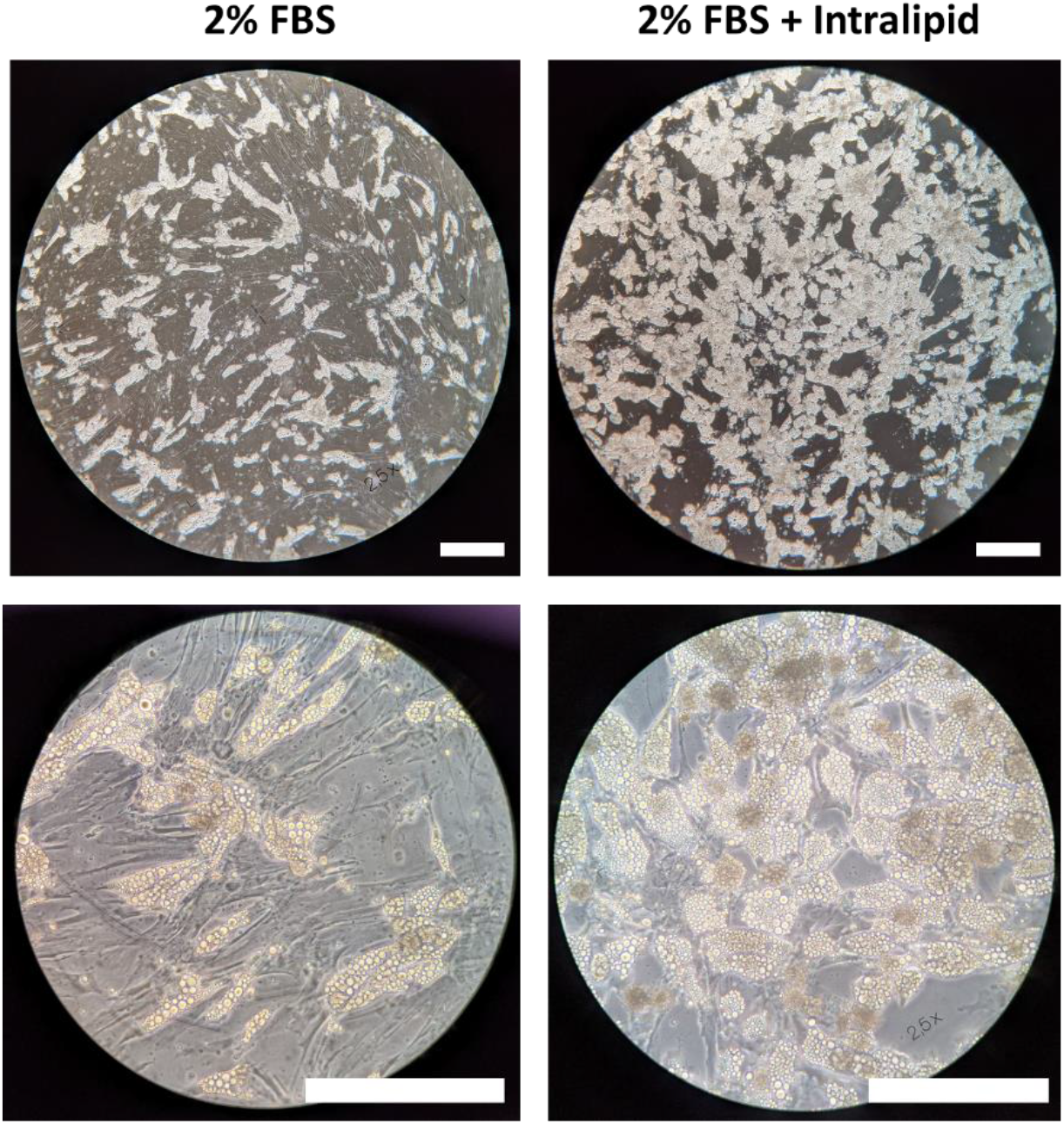
Phase contrast micrographs showing the degree of lipid accumulation in primary porcine adipocytes (DFAT cells) adipogenically differentiated with and without 500 μg/ml Intralipid for 11 days. Scale bars represent 250 μm.

**Supplemental Figure 9.**
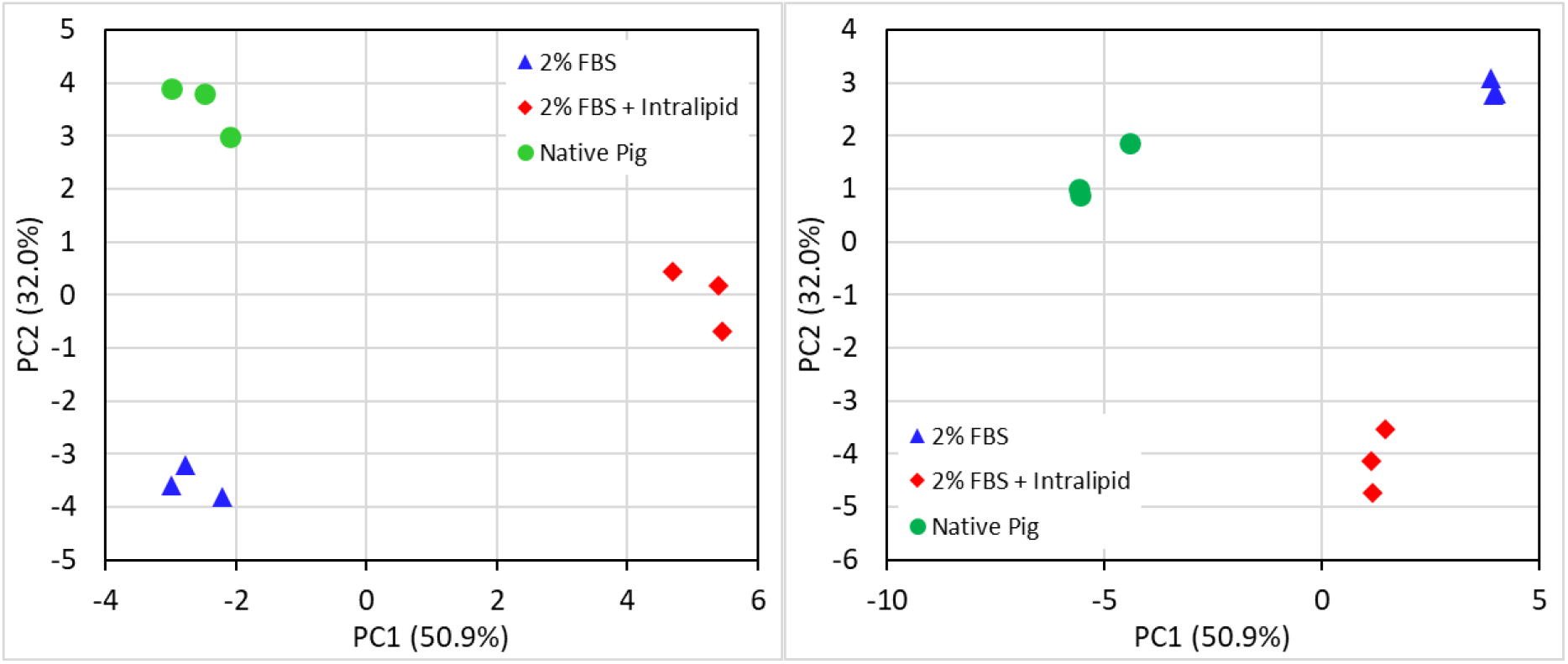
Principal component analyses of **(Left)** <u>triacylglyceride</u> (TAG) and **(Right)** phospholipid fatty acid compositions for *in vitro* and native <u>porcine</u> fats (all fatty acids). Circles (•) represent native pig fats, triangles (▴) represent *in vitro* adipocytes sans Intralipid, and diamonds (♦) represent *in vitro* adipocytes with Intralipid.

**Supplemental Figure 10.**
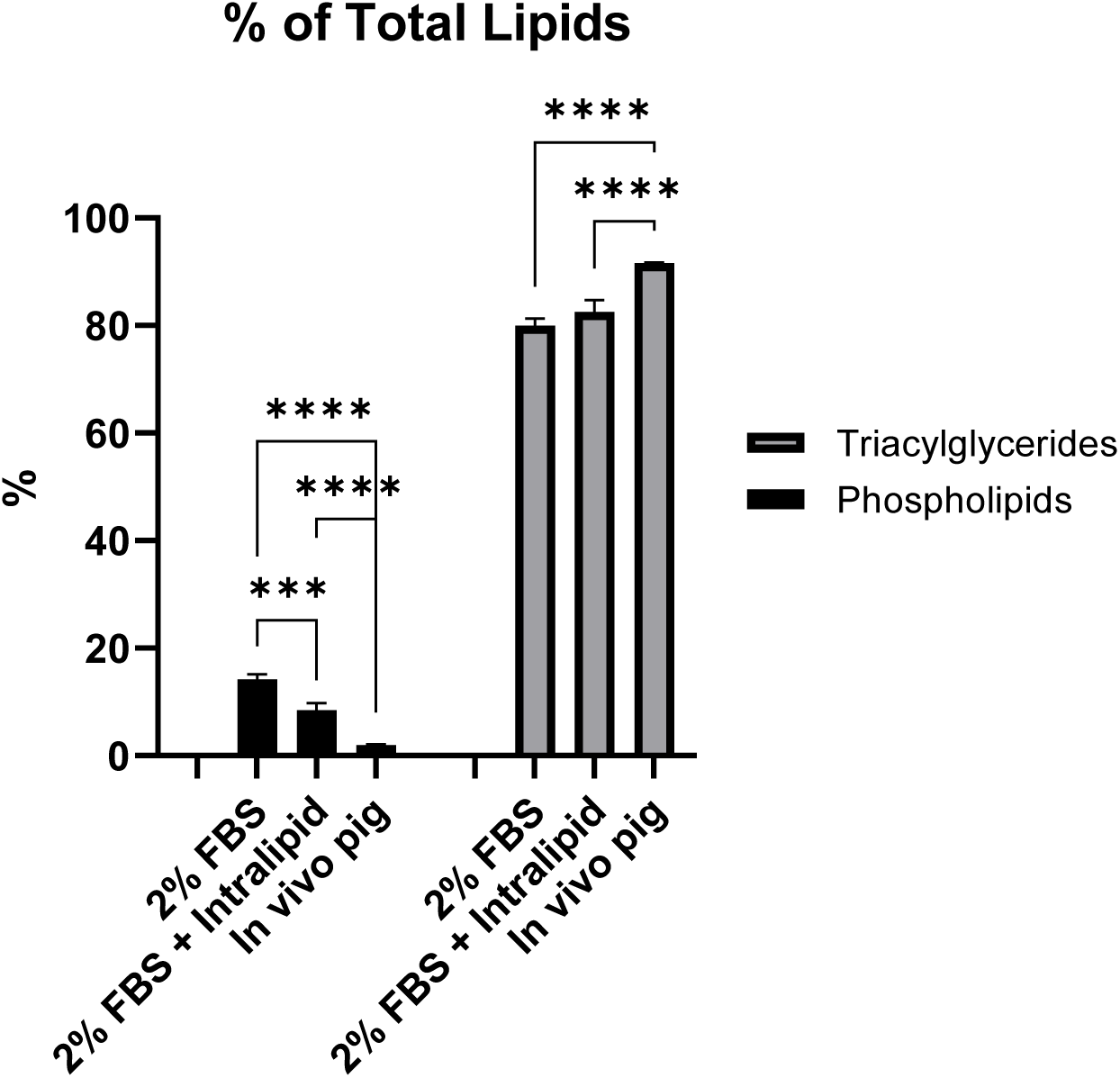
The proportion of triacylglycerides and phospholipids as a percentage of total intracellular lipids, for *in vitro* and native <u>porcine</u> fats. (***) represents p < 0.001 and (****) represents p < 0.0001.

